# Epithelial cytokeratin 6a restricts secretory autophagy of proinflammatory cytokines by interacting with Sec16A

**DOI:** 10.1101/2024.01.04.574264

**Authors:** Anand Bhushan, Jonathan K. Chan, Yan Sun, Fariba Ghiamati, Jack S Crabb, Geeng-Fu Jang, Belinda Willard, John W Crabb, Connie Tam

**Affiliations:** Department of Ophthalmic Research, Cleveland Clinic Cole Eye Institute and Lerner Research Institute, Cleveland, OH, United States; Mass Spectrometry Core, Lerner Research Institute of the Cleveland Clinic, Cleveland, OH, United States; Department of Ophthalmology, Cleveland Clinic Lerner College of Medicine of Case Western Reserve University, Cleveland, OH, United States

## Abstract

Epithelial cells form a crucial barrier against harmful microbes and inflammatory stimuli. Restraining inflammatory responses at the corneal barrier is necessary for avoiding sight-threatening tissue damage. Yet, epithelial cell-intrinsic mechanisms that dampen inflammation are largely unexplored. Keratin 6a (K6a) is a common type II cytokeratin highly expressed in corneal and other stratified epithelial cells. In a mouse model of sterile corneal inflammation, K6a knockout mice exhibit disease exacerbation. Here, we investigated cell-intrinsic mechanisms by which cytoplasmic K6a curbs corneal inflammation. We stimulated wild-type (WT) and K6a siRNA-knockdown (K6a-KD) human corneal epithelial (hTCEpi) cells with inflammatory *P. aeruginosa* culture supernatant. Our results showed that, under both basal and inflammatory conditions, K6a-KD cells secreted higher levels of cytokines and chemokines (IL-1α, IL-6, IL-8, CXCL1, CCL20) as compared to WT cells. K6a-KD cells also had increased level of LC3-II, a marker for autophagosomes, while autophagic degradation of SQSTM1/p62 remained unchanged. In K6a-KD cells, the majority of LC3-II puncta were associated with non-acidified autophagosomes rather than acidified autolysosomes. Upon stimulation, IL-8 was found to co-localize with LC3-II by confocal microscopy. Mechanistically, mass spectrometric analysis of K6a immunoprecipitates identified Sec16A, a protein involved in secretory autophagy, as an interacting partner of K6a. Further experiments showed that knocking down key proteins involved in autophagosome formation (ATG5) and the secretory autophagy process (Sec16A, GRASP55, Rab8) abolished the augmentative effect of K6a-KD on cytokine and chemokine secretion. These findings reveal a novel repressive role of K6a in secretory autophagy-mediated proinflammatory cytokine secretion and provide new insights into cell-intrinsic mechanisms of inflammation control at epithelial barriers.

## INTRODUCTION

Microbial keratitis, known as infectious corneal inflammation, poses a significant challenge to worldwide eye health. Although inflammation serves as a natural defense against microbial invaders, the influx of immune cells during an infection can result in substantial damage to the eye’s tissues. Typical risk factors for microbial keratitis involve contact lens wear, corneal trauma, insufficient tear production, immunosuppression, and systemic diseases such as diabetes ^1–3^. Most often, medical treatment incorporating a range of antimicrobial therapies holds great promise in effectively treating the disease. However, there are situations where treatment proves ineffective, resulting in serious consequences like partial or total vision loss, or even the loss of the eye itself ^4, 5^. The observed pathology of microbial keratitis results from the interplay between infecting pathogens and host immune response. The host’s reactions, which take place after the disease has started (such as infiltration of immune cells and the activation of adaptive immunity), are being extensively studied with the aim to devise novel therapies for managing inflammation-associated damages ^6–8^. In addition to immune cells, epithelial cells serve as immune sentinels, orchestrating immune responses through the release of cytokines, antimicrobial peptides, and the expression of MHCII proteins ^9–11^. The corneal epithelium, being the outermost layer of the cornea, serves as the initial barrier against external threats. The corneal epithelium’s resistance to microbial threats can be attributed to extrinsic and intrinsic factors like the expression of diverse junction molecules, the mucin layer on the apical surface of the corneal epithelium, and the production of various antimicrobial peptides ^12–16^.

Cytokeratins are the predominant and highly conserved intermediate filament proteins in epithelial cells. The expression of keratin 6a (K6a), a type II cytokeratin, is observed in various stratified epithelial cells, such as those in the skin, oral cavity, genital tract, and cornea ^17, 18^. Of note, mutation of K6a is associated with Pachyonychia Congenita, an autosomal dominant genodermatosis, and impaired mitochondrial quality control ^19, 20^. Recent advancements have unveiled the crucial regulatory function of epithelial keratins in processes related to inflammation and immunity; however, their precise molecular mechanisms are still unclear ^21, 22^. For instance, an upregulation of keratin 8 (K8) was found in the retina of patients with neovascular age-related macular degeneration. This upregulation appears to confer a protective function against oxidative stress-induced necrotic cell death in retinal pigment epithelial cells through the induction of autophagy ^23^. In our previous studies, we reported that corneal epithelial cells exhibit a direct antibacterial response ^16^. This response entails reorganization of the K6a filament network to boost the level of cytosolic K6a. This soluble K6a is processed by the ubiquitin-proteasome system leading to production of a set of short yet highly effective antimicrobial peptides, known as keratin-derived antimicrobial peptides (KAMPs) ^16, 24, 25^. In a mouse model of *P. aeruginosa* and *S. aureus* keratitis, we demonstrated that treatment with KAMPs not only kills bacteria in the infected corneas but also controls infiltration of neutrophils and macrophages leading to significant reduction of vision-threatening corneal opacification and tissue damage ^26^. Recently, we also reported that C57BL/6 mice with K6a gene knockout exhibit higher severity of corneal inflammation with and without infection, and that cytosolic K6a in corneal epithelial cells interacts with ELKS to suppress Toll-like receptor-mediated IKKɛ-dependent (non-canonical) NF-κB activation, thereby controls inflammatory cytokine expression^27^.

The majority of proteins containing signal peptides are destined for the cell surface or extracellular space and released through the conventional secretory pathway. Initially, these proteins enter the endoplasmic reticulum (ER) in their nascent form with the aid of signal peptide recognition sequences. Subsequently, these cargo proteins exit the ER at specific membrane regions known as ER exit sites (ERES), where vesicles coated with the cargo-containing complex known as coat protein complex II (COPII) are formed. This coat is responsible for cargo selection, either through Sec proteins or through interactions with cargo receptors. The cargo proteins are then transported to the Golgi apparatus, where they undergo modifications, processing, and sorting, and are directed to their final destination ^28–30^. On the other hand, there are proteins being delivered to the plasma membrane or extracellular space through distinct and uncommon secretion processes. These unconventional pathways enable secretion of cytoplasmic proteins that do not possess signal peptides ^30–32^, including the autophagy-dependent transport of yeast acyl coenzyme A-binding protein (Acb1) and mammalian pro-inflammatory cytokines (IL-1β, IL-18 and IL-33) and alarmin (HMGB1) ^33–38^. Notably, this unique unconventional protein secretion mechanism (also known as secretory autophagy) is not limited to leaderless proteins ^39–41^.

Increasing attention to the role of secretory autophagy in the release of inflammatory mediators ^34, 40, 42–44^ could be attributed to its co-occurrence with unconventional secretion ^32, 45^. The understanding of processes and molecules involved in secretory autophagy is starting to unfold. This includes the involvement of caspase 1, ER-exit site protein Sec16A, the peripheral Golgi protein GRASP55, as well as small Ras GTPases Rab8, which could function as a plasma membrane tether for membrane compartments during inflammation ^34, 42, 44, 46^. Here, we report a novel mechanism by which cytosolic K6a suppresses inflammation. Starting with proteomic analysis of human K6a immunoprecipitates showing an enrichment of autophagy pathway along with protein secretion pathways, we found that cytosolic K6a interacts with a key secretory pathway protein, Sec16A, and in turn, suppresses the formation of GRASP55^+^ LC3-II^+^ secretory autophagosomes to control secretion of cytokines and chemokines. This study reveals a novel repressive role of K6a in secretory autophagy-mediated proinflammatory cytokine secretion and provides new insights into cell-intrinsic mechanisms of inflammation control at epithelial barriers.

## MATERIALS AND METHODS

### Cell culture

Human telomerase corneal epithelial (hTCEpi) cells ^47^ were cultured in KGM-2 media (Lonza) without antibiotics. Cells were maintained at 37°C in a 5% CO_2_ humidified environment, with media changed every other day. For siRNA gene silencing, 0.1 million hTCEpi cells were seeded while at the same time being transfected with 100 nM of ON-TARGETplus non-targeting control (NTC) pool siRNAs (D-001810), or human K6a-, ATG5-, Sec16A, GRASP55-, Rab8a-, Rab8b-specific SMARTpool siRNAs (L-012116; L-004374; L-026032; L-019045; L-003905; L-008744; Horizon Discovery) using Lipofectamine RNAiMAX reagent (Invitrogen). Media was refreshed 24 h post-transfection, and the cells were incubated for an additional 48 h. Subsequently, cells were subjected to stimulation with 20% (v/v) bacterial culture supernatant or 20% (v/v) tryptic soy broth (TSB) in KGM-2 media. To prepare sterile stock solutions of bacterial culture supernatant and broth control, 20 ml TSB with or without *Pseudomonas aeruginosa* PAO1 inoculation was incubated in a shaker at 37°C and 180 rpm for 18 h, followed by centrifugation (3000 RCF, 20 min) to remove bacteria and then filter sterilization. After 18-20 h stimulation, hTCEpi cell culture supernatants were collected for cytokine measurement, and cells were processed for immunofluorescence microscopy or immunoblotting.

### Cytokine measurement

Cytokines and chemokines in cell culture supernatants were measured using enzyme-linked immunosorbent assay (ELISA) (R&D Systems) with four-parameter logistic (4-PL) fitted standard curves according to manufacturer’s protocol. Multiplex assays were performed using the Human Cytokine/Chemokine Panel A 48-plex Discovery Assay (service provided by Eve Technologies).

### Generation of stable expression hTCEpi cells

The human K6a sequence flanked by an amino-terminal HA tag and a carboxyl-terminal Flag tag (HA-hK6a-FLAG) was generated by PCR using pUC57-hK6a as a template which is codon-optimized for mammalian cell expression (Genscript). The sequence was inserted into lentiviral expression vector pCDH533-IRES-Neo (System Biosciences) to become pCDH533-HA-hK6a-Flag-IRES-Neo. To produce lentiviral particles, 7 μg of lentiviral vectors was mixed with a Lenti-X packaging (VSV-G) single shots tube (Clontech) in a final volume of 600 μl for 15 min before the transfection mix was added to subconfluent (80-90%) Lenti-X 293T cells (Clontech) grown on a 10-cm tissue culture dish. The medium containing the virus particles was collected and pooled at 48 and 72 hours after transfection. After removal of cell debris by centrifugation at 500 RCF for 15 min, clarified supernatant was combined with Lenti-X concentrator (Clontech) at a 3-to-1 volume ratio. After overnight incubation at 4°C, the lentivirus particles were pelleted by centrifugation at 1,200 RCF for 45 min at 4°C and resuspended in 500 μL of KGM-2 media. The virus titer was determined using Lenti-X GoStix Plus (Clontech). To transduce hTCEpi cells, 0.1 million cells/well seeded on 24-well plates were incubated with lentiviral particles encoding pCDH533-IRES-Neo or pCDH533-HA-K6a-Flag-IRES-Neo at 40 multiplicities of infection in the presence of 2 μg/ml polybrene in KGM-2 media for 2 h, followed by two washes with KGM-2. Cells were propagated for 1 week before they were selected for stable expression of neomycin-resistant gene in the presence of G418 at 750 μg/ml.

### Immunoprecipitation

hTCEpi cells stably expressing pCDH533-IRES-Neo or pCDH533-HA-K6a-Flag-IRES-Neo vectors in T175 flasks were lysed on ice for 15 min using cell lysis buffer [1% (v/v) Triton X-100 in 50 mM Tris, pH 7.2, 150 mM NaCl, 1 mM EDTA, 1× Halt protease inhibitor cocktail (Thermo Scientific) and 1 mM PMSF], followed by cell debris removal by centrifugation at 15,000 RCF for 15 min at 4 °C. For each immunoprecipitation, 2.5 mg lysate proteins were mixed with 200 µL of anti-Flag M2 (Sigma-Aldrich) or anti-HA (Thermo Scientific) magnetic beads overnight at 4°C with rotation. After removal of unbound materials by three washes in lysis buffer and equilibration in TBS buffer (50 mM Tris, pH 7.2, and 150 mM NaCl), the bound proteins were eluted from the beads by incubation in 500 μl of free Flag peptides at 150 μg/ml in TBS, 1× Halt protease inhibitor cocktail and 1 mM PMSF for 30 min at 4 °C. The eluate was concentrated 10-fold to 50 μL using Vivaspin 500 centrifugal concentrator (Sartorius) with molecular weight cut-off at 10 kDa.

### Mass spectrometry and pathway analysis

Anti-Flag antibody precipitated proteins in the concentrated eluate were resolved in a 4-15% Criterion TGX gel (BioRad) and stained by Colloidal Blue dye (Invitrogen). Ten gel areas within the region >64 kDa were excised and digested with trypsin for peptide extraction. Samples were analyzed by LC-MS/MS using an Orbitrap Fusion Lumos Tribrid mass spectrometer (Thermo Scientific) equipped with a Dionex Ultimate 3000 nano UHPLC system, and a Dionex (25 cm x 75 µm id) Acclaim Pepmap C_18_, 2-μm, 100-Å reversed-phase capillary chromatography column. Peptide digests (5 μl) were injected onto the reverse phase column and eluted at a flow rate of 0.3 μl/min using mobile phase A (0.1% formic acid in H_2_O) and B (0.1% formic acid in acetonitrile). The gradient was held at 2% B for 5 min, increased linearly to 45% B in 90 min, then 90% B in 15 min, and maintained at 90% B for 5 min. The samples were analyzed using a data-dependent acquisition method which involved full MS1 scans from 300-1700 Da in the Orbitrap MS at a resolution of 120000. This was followed by HCD (0.7 Da isolation window) on precursors with charge states of 2 to 7 at 32% NCE with orbitrap detection at a resolution of 50000. MS/MS spectra were acquired for 3 seconds. Dynamic exclusion was enabled where ions within 10 ppm were excluded for 60 seconds. The Mascot 2.7.0 search engine and the UniProt human database version 2021_03 (20,387 sequences) were used for protein identification. The database search parameters were restricted to three missed tryptic cleavage sites, a precursor ion mass tolerance of 10 ppm, a fragment ion mass tolerance of 20 ppm and a false discovery rate (FDR) of ≤ 1%. Protein identification required the detection of at least two unique peptides per protein and a Mascot ion score ≥ 25. Spectral counting normalized to the total amino acid length was used to estimate relative protein abundances in a given gel band. Pathway analysis was conducted by REACTOME (https://reactome.org/PathwayBrowser/#TOOL=AT) ^48^. UniProt accession IDs of K6a co-immunoprecipitated proteins as identified by LC-MS/MS were used as input data. The options of converting non-human identifiers in the input data to human equivalents and expanding REACTOME pathways to include protein-protein interactors from the IntAct database were used.

### Immunoblotting

Cells were lysed directly on the plate using ice-cold RIPA buffer [25 mM Tris (pH 7.4), 150 mM NaCl, 1% NP40, 0.1% SDS, and 1% Sodium Deoxycholate]. Protein concentration of each sample was determined using BCA kit (Thermo Scientific). Whole lysate samples containing 15-50 μg protein were electrophoresed in 4-15% SDS-PAGE gel (BioRad) and transferred onto nitrocellulose membranes using Trans-Blot Turbo Transfer System (Bio-Rad) or by wet transfer (specifically K6a and sec16A blots). Membranes was then blocked with 5% nonfat dry milk powder in TBST, followed by primary antibodies – rabbit polyclonal anti-LC3 [GTX127375] (GeneTex) (1:1000), mouse monoclonal anti-p62 [GT1478] (Invitrogen) (1:2000), rabbit anti-human K6a serum (custom made by New England Peptide) (1:1000), rabbit polyclonal anti-Sec16A (cat. no. A300-648A, Bethyl Laboratories) (1:1000), mouse monoclonal anti-GORASP2/GRASP55 [1C9A3] (Proteintech) (1:2000), and mouse monoclonal anti-beta-actin [8H10D10] (Cell Signaling) (1:1000). HRP-conjugated goat anti-rabbit IgG or goat anti-mouse IgG were used as secondary antibodies (BioRad). Western blot images were scanned, and band intensities were quantified using ImageJ.

### Confocal immunofluorescence microscopy

Cells were cultured and stimulated on chamber slides with 4 or 8 wells (EMD Millipore). Following the experiment, cells were fixed by immersing into 4% paraformaldehyde (PFA) in PBS for 30 min. Fixed cells were rinsed twice with PBS, followed by permeabilization with 0.1% Triton X-100 in PBS for 15 min. Subsequently, cells were blocked with 10% goat serum in PBS for at least an hour. For immunostaining, rabbit polyclonal anti-LC3 [GTX127375] (GeneTex) (1:250), mouse monoclonal anti-IL-8 [O-IL8-15] (Abcam) (1:100), rabbit polyclonal anti-Sec16A (cat. no. A300-648A, Bethyl Laboratories) (1:250), and mouse monoclonal anti-GORASP2/GRASP55 [1C9A3] (Proteintech) (1:500) were used to incubate cells for 2-4 hours at room temperature. Cells were then subjected to triple washes with PBST, then treated with secondary antibodies conjugated with Alexa fluor 488 or Alexa fluor 555 (1:500). ProLong Gold Antifade mountant containing DNA stain DAPI was used as the mounting medium. Cells were observed using Leica SP8 confocal microscope. Immunofluorescence intensities, co-localization, and puncta counts were analyzed using ImageJ.

### Transmission electron microscopy (TEM)

WT and K6a-KD hTCEpi cells cultured on chamber slides were treated with 20% TSB or 20% PAO1 culture supernatant. Samples were fixed on chamber slides using a mixture of 2.5% glutaraldehyde and 4% paraformaldehyde in 0.2 M cacodylate buffer at 4°C overnight, followed by three washes with sodium cacodylate buffer (0.2M, pH 7.3) for 5 min each. Samples were post-fixed with 1% Osmium Tetroxide in water at 4°C for 60 min, followed by two washes with sodium cacodylate buffer for 5 min each, and a rinse with 2% Maleate buffer (pH 5.1) for 5 min. Samples were stained with 1% uranyl acetate in Maleate buffer for 60 min at room temperature, followed by three washes with maleate buffer for 5 min each. Sample dehydration was conducted with cold ethanol at 30%, 50%, 75%, and 95% for 5 min each, followed by three rounds of 100% ethanol at room temperature, each for 10 min. The samples were then infiltrated by replacing ethanol with 1:1 mixture of 100% ethanol and Eponate 12 medium, left overnight at room temperature, and subsequently with pure Eponate 12 medium for 4-6 hours at room temperature. Finally, the samples were embedded in pure Eponate 12 in a rubber mold and polymerized in an oven at 62°C for 24 h. Ultra-thin sections of 85 nm were cut using a diamond knife, stained with 10% uranyl acetate and lead citrate, and then observed under a Tecnai G2 SpiritBT electron microscope operating at 80 kV.

### Immuno-TEM

Experiments were performed in a 10-cm cell culture dish. At the end of the experiment, cells were collected and fixed overnight at 4°C in a solution containing 4% PFA and 0.05% Glutaraldehyde. This was followed by a dehydration step using TEOH, progressing from 30% to 100% concentrations, with each step lasting 10 min at room temperature. The infiltration process included immersing the samples first in 1:1 mixture of 100% TEOH and LR white resin overnight, followed by full immersion in 100% LR white resin for 6 hours. The embedding process was completed by allowing LR white to polymerize at 50°C. Subsequently, ultra-thin sections of 90 nm were cut using a diamond knife and mounted on nickel grids coated with formvar. For the immuno-staining protocol, the grids with the mounted sections were washed three times with PBS for 5 min each. They were then blocked with 1% BSA in PBS for 30 min at room temperature. Rabbit polyclonal anti-LC3 [GTX127375] (GeneTex) and mouse monoclonal anti-GORASP2/GRASP55 [1C9A3] (Proteintech) were diluted at a ratio of 1:100 and incubated overnight at 4°C in PBS with 1% BSA. Afterward, the grids were washed with PBS three times, each for 5 min. Following this, 0.1% BSA solution in PBS was used for a 10-min blocking step at room temperature. The secondary antibodies, 5 nm Gold-conjugate anti-rabbit IgG, and 15 nm Gold-conjugate anti-mouse were diluted at a ratio of 1:10 in 0.1% BSA in PBS and applied to the grids. Each grid was then placed on a drop of GAM (30 ul) and left for 60 min at room temperature. Subsequently, the grids were washed with PBS three times, each for 5 min, and the antibodies were fixed with 1% Glutaraldehyde in PBS for 10 min. After another round of washing with water (three times for 5 min each), the grids were stained with UA and lead citrate. Finally, the grids were dried and observed using a Tecnai G2 SpiritBT electron microscope operated at 80 kV.

### Statistical analysis

GraphPad Prism (version 10) was used to perform statistical tests. Normal distribution of data and equality of variances were determined by Shapiro-Wilk test and Brown-Forsythe test respectively. Outliers were detected by Grubb’s test. For normally distributed data, statistical significance was determined using the two-tailed Student’s t-test, one-way ANOVA with Sidak’s multiple comparisons test, or Welch ANOVA with Dunnett’s T3 multiple comparisons test. For non-normal data, Mann-Whitney U test, Wilcoxon matched-pairs test, or Kruskal-Wallis with Dunn’s multiple comparisons test was used. At least three independent repeats for each experiment had been carried out. *P* < 0.05 was considered statistically significant.

## RESULTS

### Endogenous K6a suppresses proinflammatory cytokine and chemokine release from human corneal epithelial cells

To investigate the role of K6a in the secretion of cytokines, we utilized human telomerase-immortalized corneal epithelial (hTCEpi) cells as an in vitro model. K6a was knocked down (K6a-KD) using siRNA, and we induced sterile inflammation by treating the cells with 20% *P. aeruginosa* PAO1 culture supernatant in KGM-2 media with 20% TSB as baseline control. Cell culture supernatant was collected and subjected to multiplex human cytokine assay (a panel of 48 cytokines). Among twenty-one secreted molecules detectable in the culture media, K6a-KD cells under basal conditions secreted higher levels of IL-1α, IL-6, IL-8, IL-22, IFN-γ, G-CSF, CXCL1, CXCL10, CCL5, PDGF-AB/AA and VEGF-A as compared to non-targeting siRNA treated (wild type) cells (Figure 1A). Similarly, upon stimulation by 20% PAO1 culture supernatant, K6a-KD cells also secreted higher levels of IL-1α, IL-6, IL-8, GROα/CXCL-1 as compared to wild type cells (Figure 1A). Next, we validated some of these findings by traditional ELISA (Figure 1B), which was in line with our recent findings ^27^. These results indicated that K6a in corneal epithelial cells negatively regulates secreted levels of proinflammatory mediators.

**Figure 1.**
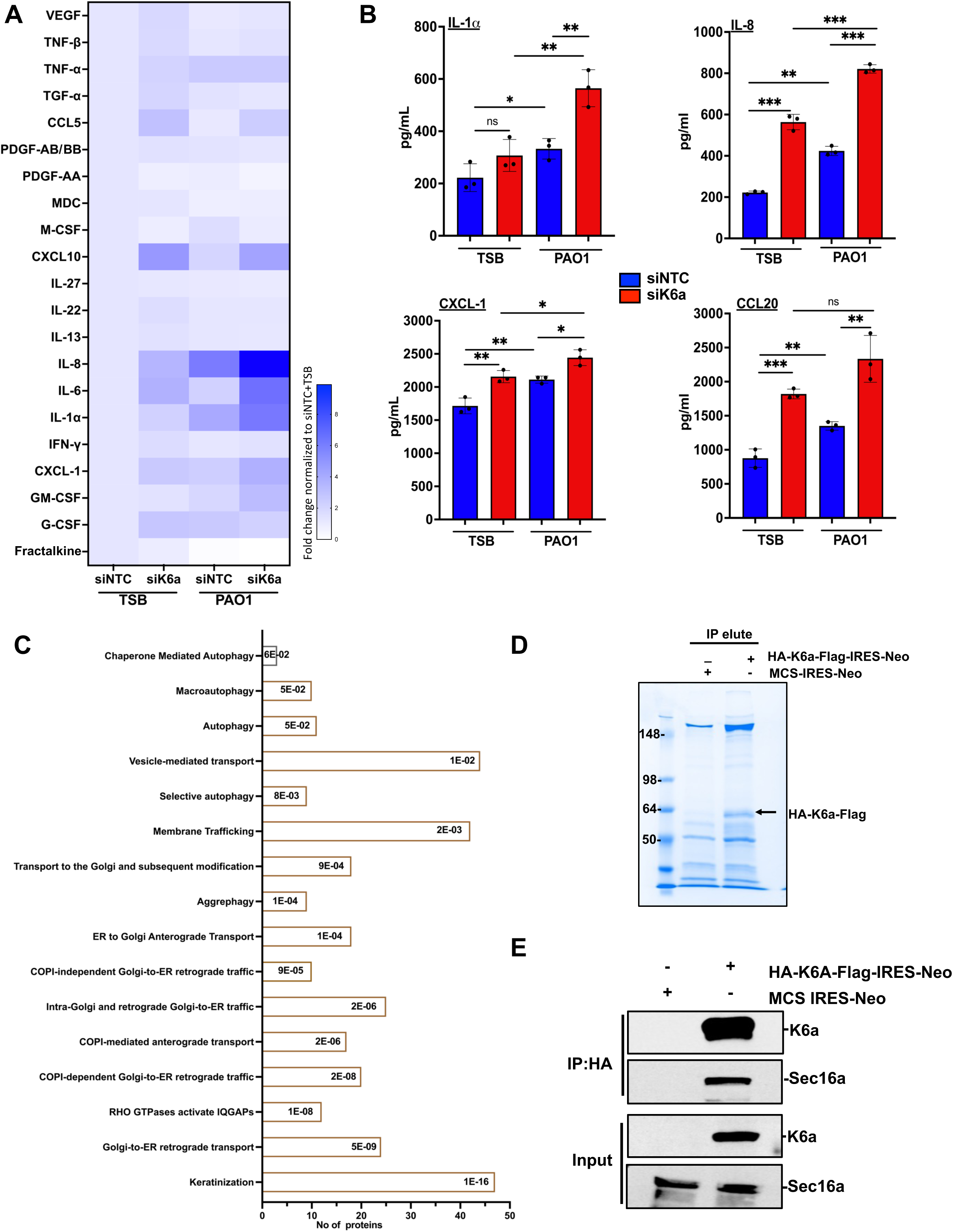
K6a in human corneal epithelial cells (hTCEpi) controls secretion of cytokines and chemokines, participates in autophagy and protein secretory pathways, and interacts with Sec16A. (**A-B**) hTCEpi cells transfected with non-targeting siRNA (siNTC) and K6a-specific siRNA (siK6a) were exposed to 20% sterile *P. aeruginosa* PAO1 culture supernatant or 20% tryptic soy broth (TSB) in KGM-2 media for 18-20 h. Cell culture supernatants were collected for cytokine and chemokine quantification (n = 3 independent experiments). (**A**) Multiplex assay was performed; data normalized to baseline control (siNTC+TSB) is represented by a heat map with gradient color bars showing fold changes. (**B**) ELISA of IL-1α, IL-8, CXCL1 and CCL20 (mean ± SD). Statistical significance was determined by a two-tailed unpaired t-test. *P < 0.05, **P < 0.01, ***P < 0.001; ns, P > 0.05. (**C**) Anti-Flag immunoprecipitation of whole-cell lysate from hTCEpi cells expressing HA-K6a-Flag followed by LC-MS/MS and pathway analysis of identified proteins. Bar graph representation shows each enriched pathway with the number of identified proteins and p-value. (**D-E**) LC-MS/MS identification of Sec16A was verified by anti-HA immunoprecipitation of whole-cell lysate from hTCEpi cells expressing HA-K6a-Flag. (**D**) Eluted proteins were analyzed by SDS-PAGE and Colloidal Blue staining. The arrow indicates the band of HA-K6a-Flag. (**E**) Immunoblotting of eluted proteins confirmed Sec16A as an interacting protein of K6a.

Next, we conducted a proteomic study to uncover the underlying mechanisms by which endogenous K6a in corneal epithelial cells controls inflammation, especially through regulation of proinflammatory cytokine and chemokine secretion. First, we generated hTCEpi cells that stably expressed HA-K6a-FLAG and performed K6a immunoprecipitation with anti-FLAG antibody, followed by mass spectrometry (MS) analysis on eluted proteins that were larger than 65 kDa. We effectively identified 642 proteins (each with a minimum of 2 peptide counts), including endoplasmic reticulum (ER) export mediator Sec16A and autophagy-related protein ATG16L, co-precipitated with HA-K6a-FLAG. All protein hits were then subjected to pathway analysis using the publicly available database of biological pathways ‘Reactome’. Our analysis revealed several enriched pathways, notably including autophagy-related pathways and protein secretion pathways (Figure 1C). It is established that autophagy plays a crucial role in regulating various aspects of the immune response, thus impacting inflammatory processes ^49^. Among the immunoprecipitated proteins identified by mass spectrometry, we focused on Sec16A and verified its interaction by performing HA-K6a-FLAG immunoprecipitation again with anti-HA antibody (Figure 1D) followed by immunoblotting of sec16A (Figure 1E). Sec16A is an essential protein involved in the process of protein secretion in eukaryotic cells. It is known to play a crucial role in the early stages of the secretory pathway, specifically the endoplasmic reticulum (ER) exit site (ERES) organization and vesicle formation ^50–52^. Apart from its role in conventional protein secretion, Sec16A is also involved in unconventional protein transport pathways including autophagosome formation ^41, 53, 54^. Our findings raise the intriguing hypothesis that both autophagy and secretory pathways are intricately involved in corneal inflammation, and their regulation is influenced by the presence of K6a. Growing evidence suggested a close connection between the secretory and autophagy pathways, with some regulatory components being common to both pathways ^32, 43, 45, 49^.

### K6a restricts non-degradative autophagosome biogenesis involved in the secretion of IL-8

To examine potential impacts of K6a on the level of autophagy in hTCEpi cells, we utilized siRNA to knock down K6a expression and then assessed the number of autophagosomes using confocal immunofluorescence microscopy ^55–57^. When we compared the levels of LC3-II puncta under basal condition (20% TSB treatment), increased puncta per cell was observed in K6a-KD cells as compared to WT cells (Figure 2A). To induce inflammation, we treated both cells with 20% PAO1 culture supernatant. Again, K6a-KD cells exhibited a substantial increase in LC3-II puncta per cell when compared to WT cells (Figure 2A), consistent with the results observed under the basal condition. Using transmission electron microscopy (TEM) to observe the ultrastructure of hTCEpi cells ^58^, a higher number of double-membrane autophagic vesicles was found in K6a-KD cells than WT cells (Figure S1). In addition, we found an increase in the number of LC3-II puncta when K6a-KD cells were stimulated with the bacterial culture supernatant (Figure 2A).

**Figure 2.**
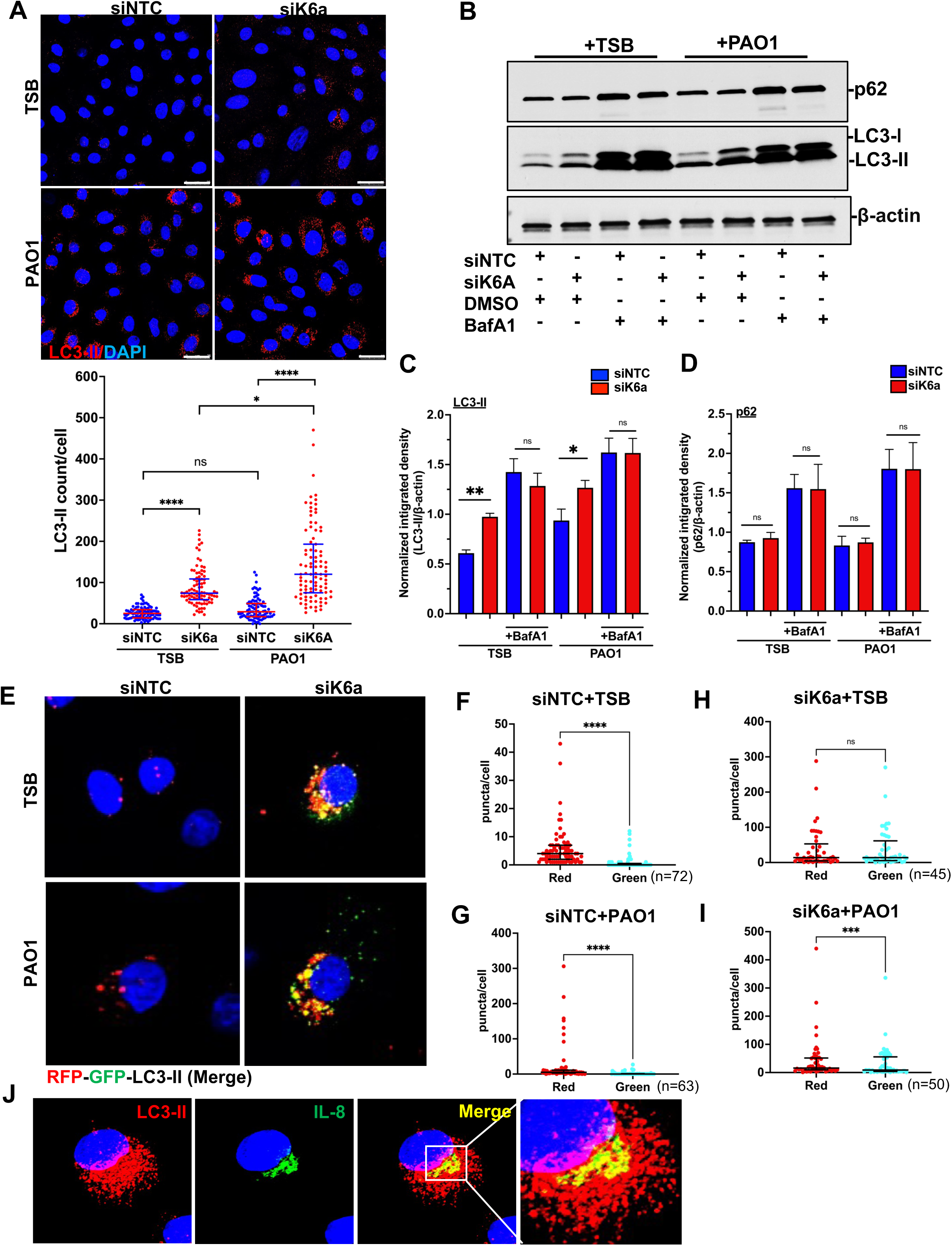
Knockdown of K6a in hTCEpi cells promotes non-degradative autophagosome formation. (A-D) hTCEpi cells transfected with non-targeting siRNA (siNTC) and K6a-specific siRNA (siK6a) were exposed to 20% sterile *P. aeruginosa* PAO1 culture supernatant or 20% tryptic soy broth (TSB) for 18-20 h. **(A)** Representative images of confocal immunofluorescence microscopy using anti-LC3 antibody. scale bar = 25µm. LC3-II puncta/cell was quantified by ImageJ. Data (median with interquartile range) was pooled from 3 independent experiments with 2-3 fields per experiment and 10-20 cells per field (totaling 88-95 cells per group). The Kruskal-Wallis with Dunn’s multiple comparisons test was used for statistical analysis. **(B-D)** hTCEpi cells were stimulated in the presence of Bafilomycin A (BafA1, 100nM) or DMSO (vehicle control). **(B)** The levels of LC3 and p62 were detected by immunoblotting. Integrated densities of **(C)** LC3-II and **(D)** p62 protein bands were quantified by ImageJ and normalized to β-actin. Data is shown as mean ± SD from three independent experiments (n=3). Two-tailed unpaired t-test was used for statistical analysis. **(E-I)** hTCEpi cells were subjected to transfection with NTC and K6a siRNA, followed by transduction with baculovirus carrying RFP-GFP-LC3-II tandem probe at a multiplicity of infection (MOI) of 1:80 for 6 hours. Subsequently, cells were treated with 20% TSB or 20% sterile PAO1 culture supernatant for 24 h before imaging by confocal microscopy. RFP (red) and GFP (green) puncta of each cell were quantified by ImageJ. Data was pooled from two independent experiments and represented as median and interquartile range. Statistical significance was determined by Wilcoxon matched-pairs test. (**J)** Co-localization of LC3-II and IL-8. K6a was knocked down by siRNA in hTCEpi cells and stimulated with 20% sterile PAO1 culture supernatant for 20 h. Cell were fixed and stained with anti-LC3 and IL-8 antibodies. *P < 0.05, **P < 0.01, ***P < 0.001, ****P < 0.0001, ns: P > 0.05.

Since accumulation of LC3-positive vesicles can result from increased biogenesis of autophagosomes and/or decreased lysosomal degradation (autophagic flux), we compared the amount of LC3-II in lysates from WT and K6a-KD cells either untreated or treated with bafilomycin A1 (BafA1; an antagonist of vacuolar H^+^ ATPase that prevents lysosomal fusion/acidification and thus autophagic degradation) ^57, 59–61^. In agreement with the results obtained from confocal immunofluorescence microscopy and transmission electron microscopy, immunoblotting of lysates from K6a-KD cells cultured in the absence of BafA1 showed significantly elevated levels of LC3-II compared with WT cells under basal and inflammatory conditions (Figure 2B, lanes 1 and 2, 5 and 6, respectively; quantification in Figure 2C). However, in the presence of BafA1, LC3-II amount was increased to a lesser extent in K6a-KD cells (Figure 2B, lanes 1 and 3, 2 and 4 under basal condition; lanes 5 and 7, 6 and 8 under inflammatory condition; quantification in Figure 2C), arguing that knockdown of K6a might impair autophagic degradation, which contributed to a higher number of LC3-II^+^ autophagosomes.

To verify if autophagic degradation is truly impacted in K6a-KD cells, we measured the level of p62/sequestosome 1 (SQSTM1), which binds to LC3-II and serves as a substrate degraded by autophagy. When autophagic degradation is suppressed, p62 accumulates, whereas induction of degradative autophagy leads to a decrease in its levels. This characteristic renders p62 a valuable marker for examining autophagic flux ^55, 57, 62, 63^. Interestingly, in the absence of BafA1, we did not observe any difference in p62 level between K6a-KD and WT cells under both basal and inflammatory conditions (Figure 2B, lanes 1 and 2, 5 and 6, respectively; quantification in Figure 2D). Also, as the cells were treated with BafA1, we observed that p62 increased to a similar extent in both WT cells and K6a-KD cells (Figure 2B, lanes 1 and 3, 2 and 4 under basal condition; lanes 5 and 7, 6 and 8 under inflammatory condition; quantification in Figure 2D). These results ruled out the possibility that knockdown of K6a impairs autophagic degradation, and supported that increased autophagosome biogenesis in K6a-KD hTCEpi cells are not destined for lysosomal degradation, but rather for secretion.

To test this hypothesis, we introduced RFP-GFP-LC3-II baculovirus into WT and K6a-KD cells (MOI 1:80; previously published method ^34, 64^) prior to treatment with 20% TSB or PAO1 culture supernatant (Figure 2E). The assay operates on the principle that GFP (green) fluorescence is quenched under acidic conditions when autophagosomes are fused with lysosomes to become autolysosomes. In line with confocal immunofluorescence microscopy and immunoblotting of LC3-II, we observed that bacterial culture supernatant induced formation of RFP (red) puncta (with or without co-occurring green fluorescence) in WT cells (Figure 2, F-G) and K6a-KD cells (Figure 2, H-I). In addition, under both basal and inflammatory conditions, WT cells exhibited a majority of LC3-II^+^ puncta as autolysosomes with RFP fluorescence only (Figure 2, F-G), whereas a majority of the LC3-II^+^ puncta in K6a-KD cells were identified as non-acidified autophagosomes (red puncta with GFP fluorescence) (Figure 2, H-I). Taken together, the results strongly supported that K6a restricts the formation of non-degradative autophagosomes in corneal epithelial cells.

Since IL-8 was one of the cytokines upregulated in the secretion of K6a-KD cells, and several reports suggested the involvement of the autophagy pathway and its associated proteins in the secretion of IL-8 ^65, 66^, we evaluated colocalization of IL-8 and autophagosomes in K6a-KD cells stimulated with bacterial culture supernatant using confocal immunofluorescence microscopy. Measurement of Pearson’s correlation coefficient in IL-8 and LC3-II stained cells indicated a reasonably strong correlation of their co-occurrence in the same pixel (Figure 2J).

Autophagy-related proteins (ATG) shape inflammatory signaling and secretory routes for leaderless proteins ^44^. Among them, ATG5 plays a crucial role in autophagy, specifically in the elongation and maturation of the phagophore into the fully formed autophagosome ^67^. ATG5, along with other autophagy-related factors like ATG7 and BCN1, has also been shown to play an important role in the secretory autophagy pathway ^34, 39, 65, 66, 68^. To further support the notion that dysregulated biogenesis of non-degradative autophagosomes in K6a-KD hTCEpi cells contributes to uncontrolled IL-8 secretion, we employed siRNA to knock down ATG5 expression in combination with K6a. Under basal (Figure 3A) and inflammatory conditions (Figure 3B), LC3-II puncta was significantly increased in K6a-KD cells as compared to WT cells, and knockdown of ATG5 in K6a-KD cells reduced LC3-II puncta to levels similar to that of WT cells. In accordance with this observation, increased IL-8 secretion from K6a-KD cells was abolished by knocking down ATG5 simultaneously (Figure 3C). Collectively, these results demonstrated the negative regulatory role of K6a in non-degradative autophagosome biogenesis, which mediates secretion of IL-8.

**Figure 3.**
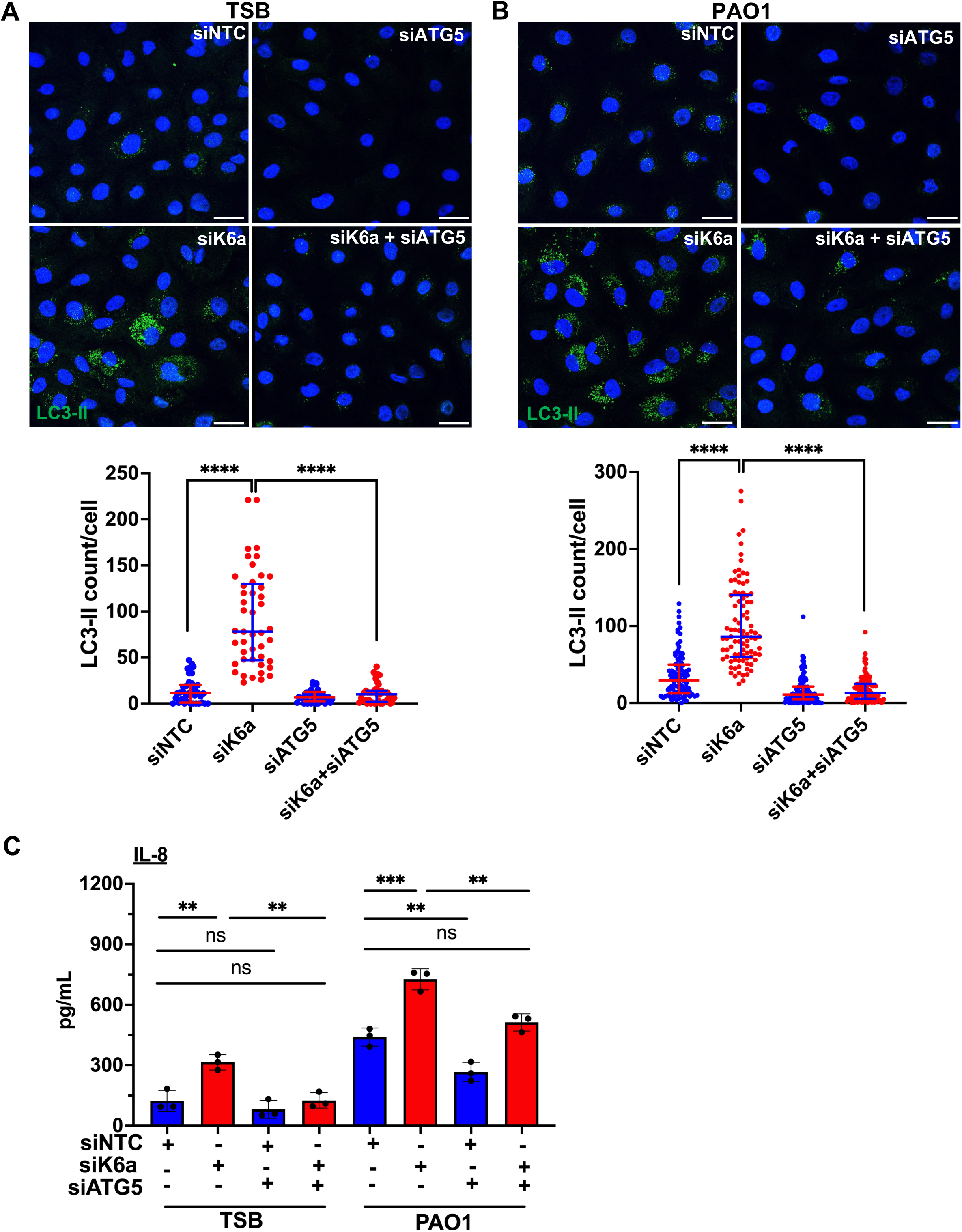
ATG5 is required for autophagy-associated IL-8 secretion from K6a-knockdown hTCEpi cells. hTCEpi cells transfected with non-targeting siRNA, or K6a– and ATG5-specific siRNA (alone or in combination), were exposed to 20% sterile *P. aeruginosa* PAO1 culture supernatant or 20% tryptic soy broth (TSB) in KGM-2 media for 18-20 h. Cells were fixed and stained with anti-LC3 antibody, and cell culture supernatants were collected for ELISA. **(A-B)** Representative images of confocal microscopy and Image J quantification of LC3-II puncta/cell. scale bar=25µm. Data (median and interquartile range) were pooled from 2-3 independent experiments, with a minimum of 2-3 fields per experiment and 10-20 cells per field. Statistical analysis was performed using Kruskal-Wallis with Dunn’s multiple comparisons test. **(C)** Secreted IL-8 in cell culture supernatants were quantified by ELISA. Data are shown as mean ± SD of three independent experiments. Statistical significance was determined by one-way ANOVA with Sidak’s multiple comparisons test. **P < 0.01, ***P < 0.001, ****P < 0.0001, ns: P > 0.05.

### K6a-Sec16A interaction highly suppresses Sec16A-dependent autophagosome formation and IL-8 secretion

In view of the interaction between K6a and Sec16A (Figure 1E), we hypothesized that knocking down K6a releases Sec16A to promote non-degradative autophagosome formation. To begin investigation of this possibility, we used confocal microscopy and observed a significant increase in the immunofluorescence signal of Sec16A in K6a-KD cells compared to WT cells under basal condition (Figure 4A) and inflammatory condition (Figure 4B), suggesting that knockdown of K6a promotes localization of Sec16A at the ER-Golgi intermediate compartment (ERGIC) and ER-exit site (ERES), where it plays a role in facilitating LC3 lipidation, a crucial process in autophagosome formation ^69, 70^. We confirmed that the protein level of Sec16A remained constant between WT and K6a-KD cells, whether the cells were treated or not with inflammatory bacterial culture supernatants (Figure 4C). Furthermore, when we knocked down Sec16A alone in hTCEpi cells under basal condition, a modest reduction of LC3-II puncta per cell was observed compared to WT cells; more importantly, knocking down both K6a and Sec16A completely reversed the major increase of LC3-II puncta in K6a-KD cells (Figure 4, D and F). A consistent observation was made when the cells were under inflammatory condition, in which knocking down Sec16A suppressed the normal and excess LC3-II puncta formation in WT and K6a-KD cells respectively (Figure 4, E and G).

**Figure 4.**
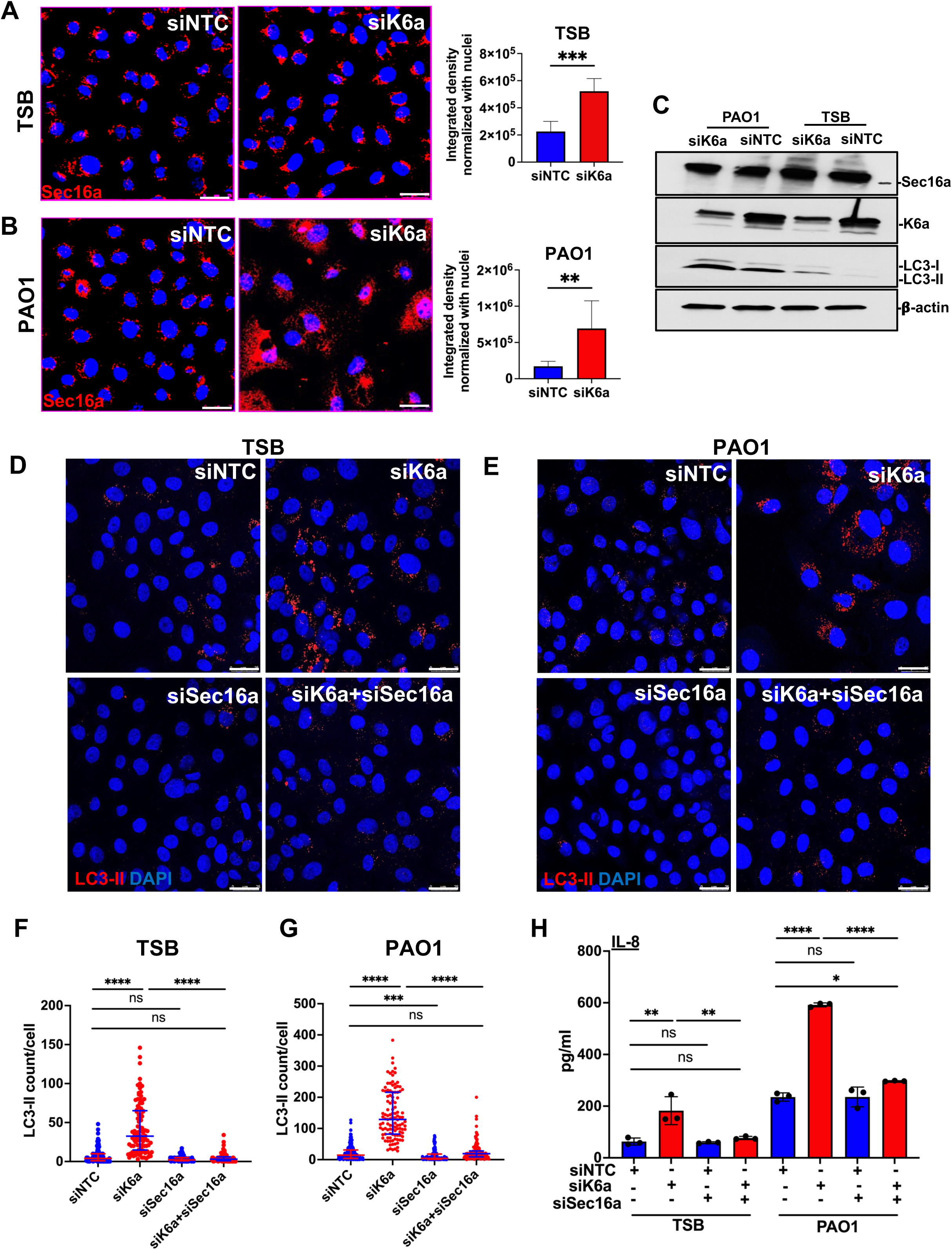
K6a restricts Sec16A-dependent autophagosome formation and IL-8 secretion. (A-C) hTCEpi cells transfected with non-targeting siRNA (siNTC) and K6a-specific siRNA (siK6a) were exposed to KGM-2 containing 20% tryptic soy broth (TSB), or 20% sterile *P. aeruginosa* PAO1 culture supernatant for 18-20 h. **(A-B)** Sec16A was immuno-stained and visualized by confocal microscopy. Scale bar=25µm. Fluorescence intensity of Sec16A was measured by ImageJ. Data (mean ± SD) from two independent experiments was analyzed by two-tailed t-test. (**C)** Western blots showed the levels of Sec16A and LC3-II in siNTC or siK6a-transfected cells under basal and inflammatory conditions. **(D-H)** hTCEpi cells were subjected to transfection with non-targeting siRNA, or K6a– and Sec16A-specific siRNA (alone or in combination) prior to stimulation with 20% TSB or 20% PAO1 for 18-20 hours. Cells were fixed and stained with anti-LC3 antibody. **(D-E)** Representative confocal microscopy images, scale bar=25µm. **(F-G)** Image J quantification of LC3-II puncta per cell. Data (median with interquartile range) were pooled from three independent experiments (n=3). For each experiment, three fields, each containing 10-20 cells, were analyzed by Kruskal-Wallis with Dunn’s multiple comparisons test. **(H)** Secreted IL-8 in cell culture supernatants were quantified by ELISA. Data is shown as mean ± SD of three independent experiments. Statistical significance was determined by one-way ANOVA with Sidak’s multiple comparisons test. *P < 0.05, **P < 0.01, ***P < 0.001, ****P < 0.0001, ns: P > 0.05.

To determine whether K6a-mediated suppression of Sec16A-dependent autophagosome formation regulates IL-8 secretion, we compared IL-8 level in culture supernatants of Sec16A-KD cells and K6a-Sec16A-double KD cells with WT cells and K6a-KD cells respectively. Although Sec16A was involved in autophagosome formation in WT cells, a change in IL-8 secretion was not found between Sec16A-KD cells and WT cells (Figure 4H). Conversely, K6a-Sec16A-double KD cells showed a significant decrease in IL-8 secretion correlating with LC3-II puncta per cell as compared to K6a-KD cells under both basal and inflammatory conditions (Figure 4H). Altogether, these results support that K6a-Sec16A interaction is a regulatory mechanism for autophagosome-mediated IL-8 release from hTCEpi cells.

### Increased autophagosome formation in K6a-KD cells and associated cytokine-chemokine secretion are dependent on GRASP55

The Golgi reassembly and stacking proteins, known as GRASPs (including GRASP55 and GRASP65 in mammals, Grh1 in yeast, and dGRASP in Drosophila), play a crucial role in autophagy-mediated unconventional secretion ^31, 34–36, 71^. According to recent reports, GRASP55 facilitates autophagy-based secretion of diverse cargo proteins through post-translational modifications as well as its redistribution to organelles like autophagosomes ^40, 72, 73^. Specifically, it has been demonstrated that GRASP55 plays a role in secretory autophagosome formation and secretion of mature IL-1β by bone marrow-derived macrophages ^34^. Given that Sec16A interacts with Golgi re-assembly and stacking protein 55 (GRASP55) to facilitate autophagosomal conveyance of mutant CFTR to the membrane surface of HeLa cells ^41^, we investigated whether Sec16A interacting partner K6a also interacts with GRASP55. Through immunoblotting of anti-HA immunoprecipitate from hTCEpi cells stably expressing FLAG-K6a-HA, we did not detect GRASP55 as an interacting protein of K6a. Nonetheless, using immunofluorescence microscopy, we noted a significantly higher signal of GRASP55 in K6a-KD cells compared to WT cells under both basal (Figure 5A) and inflammatory conditions (Figure 5B). Similar to Sec16A, we confirmed by immunoblotting that there was no discernible difference in GRASP55 protein levels between WT and K6a-KD cells, with or without inflammatory treatment (Figure 5C).

**Figure 5.**
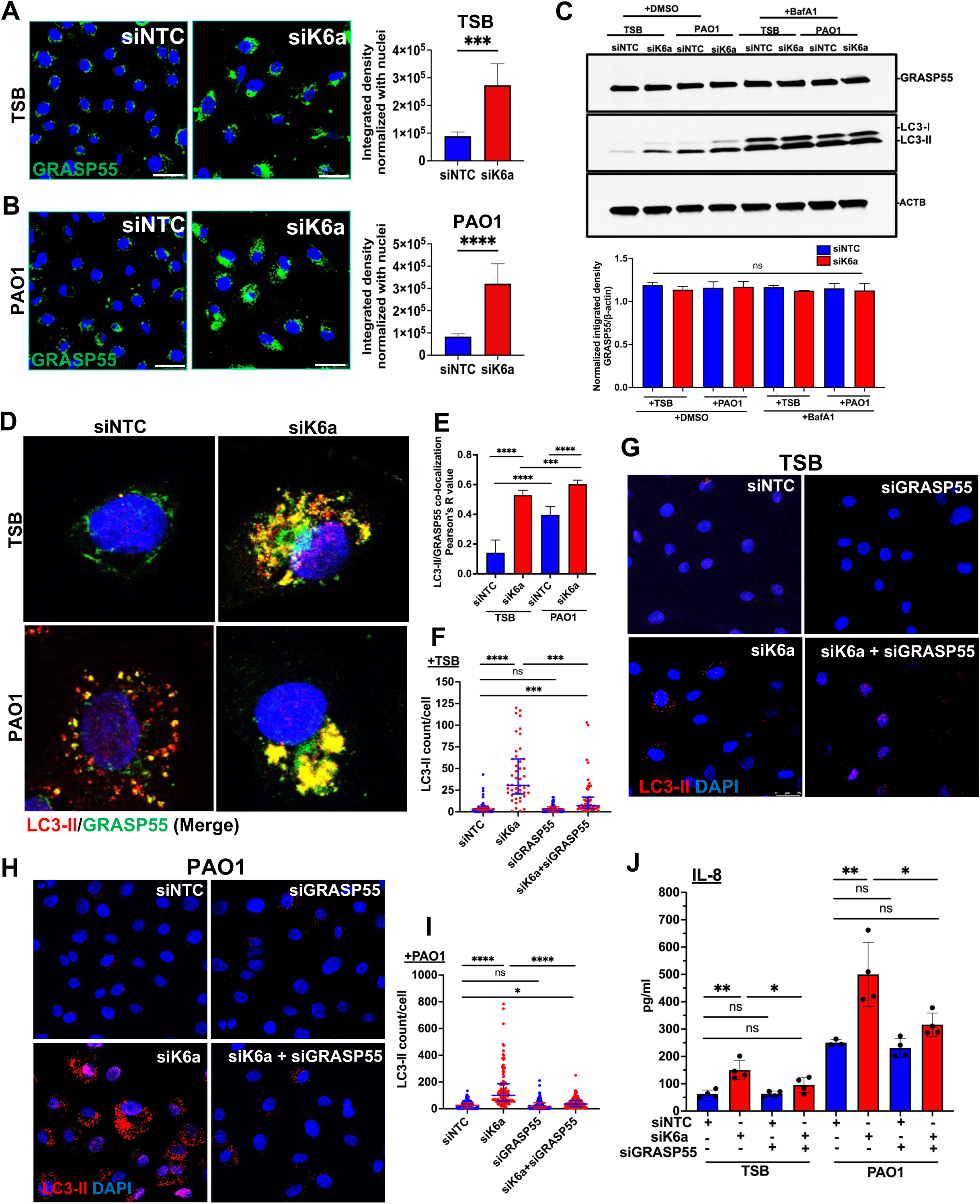
GRASP55 is required for K6a-KD-induced autophagosome formation and IL-8 secretion. **(A-E)** hTCEpi cells transfected with non-targeting siRNA (siNTC) and K6a-specific siRNA (siK6a) were exposed to KGM-2 containing 20% tryptic soy broth (TSB), or 20% sterile *P. aeruginosa* PAO1 culture supernatant for 18-20 hours. **(A-B)** GRASP55 was stained and visualized by confocal microscopy. Scale bar=25µm. Fluorescence intensity of GRASP55 was measured by ImageJ. Data (mean ± SD) from two independent experiments was analyzed by two-tailed t-test. **(C)** GRASP55 and LC3-II protein levels were detected by western blotting. GRASP55 band intensities were measured by ImageJ and normalized to β-actin. Data from two independent experiments was analyzed by one-way ANOVA. **(D-E)** Co-localization of LC3-II and GRASP55 was examined by confocal immunofluorescence microscopy. **(D)** Representative images and **(E)** ImageJ Coloc2 analysis of LC3-II and GRASP55 staining. Pearson correlation coefficients (mean ± SD) were computed from three independent experiments, and compared between groups using Welch ANOVA with Dunnett’s T3 multiple comparisons test. **(F-J)** hTCEpi cells underwent transfection with non-targeting siRNA, or K6a– and GRASP55-specific siRNA (alone or in combination) prior to stimulation with 20% TSB or 20% sterile PAO1 culture supernatant for 18-20 h. Representative confocal microscopy images of LC3-II puncta and ImageJ quantification under **(F-G)** basal condition or **(H-I)** inflammatory condition. scale bar = 25µm. LC3-II puncta/cell (median with interquartile range) was quantified by pooling data from three independent experiments with 2-3 fields per experiment and 10-20 cells per field. Statistical significance was determined by Kruskal-Wallis with Dunn’s multiple comparisons test. **(J)** Concentration of secreted IL-8 was assessed using ELISA. Data was pooled from four independent experiments and shown as mean ± SD. Statistical significance was determined by one-way ANOVA with Sidak’s multiple comparisons test. *P < 0.05, **P < 0.01, ***P < 0.001, ****P < 0.0001, ns: P > 0.05.

Drawing from our prior understanding of GRASP55’s involvement in autophagy-mediated secretion, we reasoned that increased GRASP55 immunofluorescence signal in K6a-KD hTCEpi cells might result from its recruitment to non-degradative autophagosomes. Indeed, using confocal immunofluorescence microscopy, we observed a significant increase in GRASP55 and LC3-II co-localization in K6a-KD cells compared to WT cells under basal and inflammatory conditions (Figure 5, D-E). Ultrastructural examination using immunogold transmission electron microscopy further confirmed the observed co-localization of GRASP55 and LC3-II in double-membrane vesicles (autophagosomes) of K6a-KD cells (Figure S2). As our data showed that K6a controls non-degradative autophagosome formation (Figure 2), we verified the secretory nature of GRASP55^+^LC3-II^+^ autophagosomes by inhibiting autophagic degradation with BafA1. As expected, GRASP55 protein level did not increase in the presence of BafA1 (Figure 5C). Collectively, these findings further substantiate that non-degradative GRASP55^+^ autophagosomes in K6a-KD hTCEpi cells participate in unconventional secretion processes, especially secretory autophagy, which is in line with previous reports ^34, 42^.

Next, we investigated whether GRASP55 regulates autophagosome formation and consequently, the secretion of IL-8 from hTCEpi cells. We performed knockdown experiments targeting GRASP55 alone, K6a alone, and both GRASP55 and K6a simultaneously. Again, autophagosome formation was assessed by examining LC3-II puncta per cell using confocal immunofluorescence microscopy. Compared to cells with K6a knocked down alone, those with GRASP55 and K6a double knocked down had a significant reduction in autophagosome formation under basal (Figure 5, F-G) and inflammatory conditions (Figure 5, H-I). Correlating with LC3-II puncta numbers, knocking down GRASP55 in K6a-KD cells abolished upregulation of IL-8 secretion caused by knockdown of K6a alone (Figure 5J). In contrast, no difference in IL-8 secretion was observed between WT and GRASP55-KD cells (Figure 5J). These findings suggest that GRASP55 in K6a-KD hTCEpi cells is required for secretory autophagosome formation, which in turn promotes IL-8 release.

To determine whether additional cytokine and chemokine secretions from K6a-KD cells are modulated in a GRASP55-dependent manner, we assessed the specific ones found to be upregulated by K6a knockdown in the multiplex assays (Figure 1A). As we measured the levels of IL-1α, IL-6, CCL5, CCL20, CXCL1 and CXCL10 by ELISA, we found that, similar to IL-8, increased secretion of IL-1α, IL-6, CCL20 and CXCL1 (Figure 6, A-D), but not CCL5 and CXCL10 (Figure 6, E-F), from K6a-KD cells under basal and inflammatory conditions was GRASP55-dependent.

**Figure 6.**
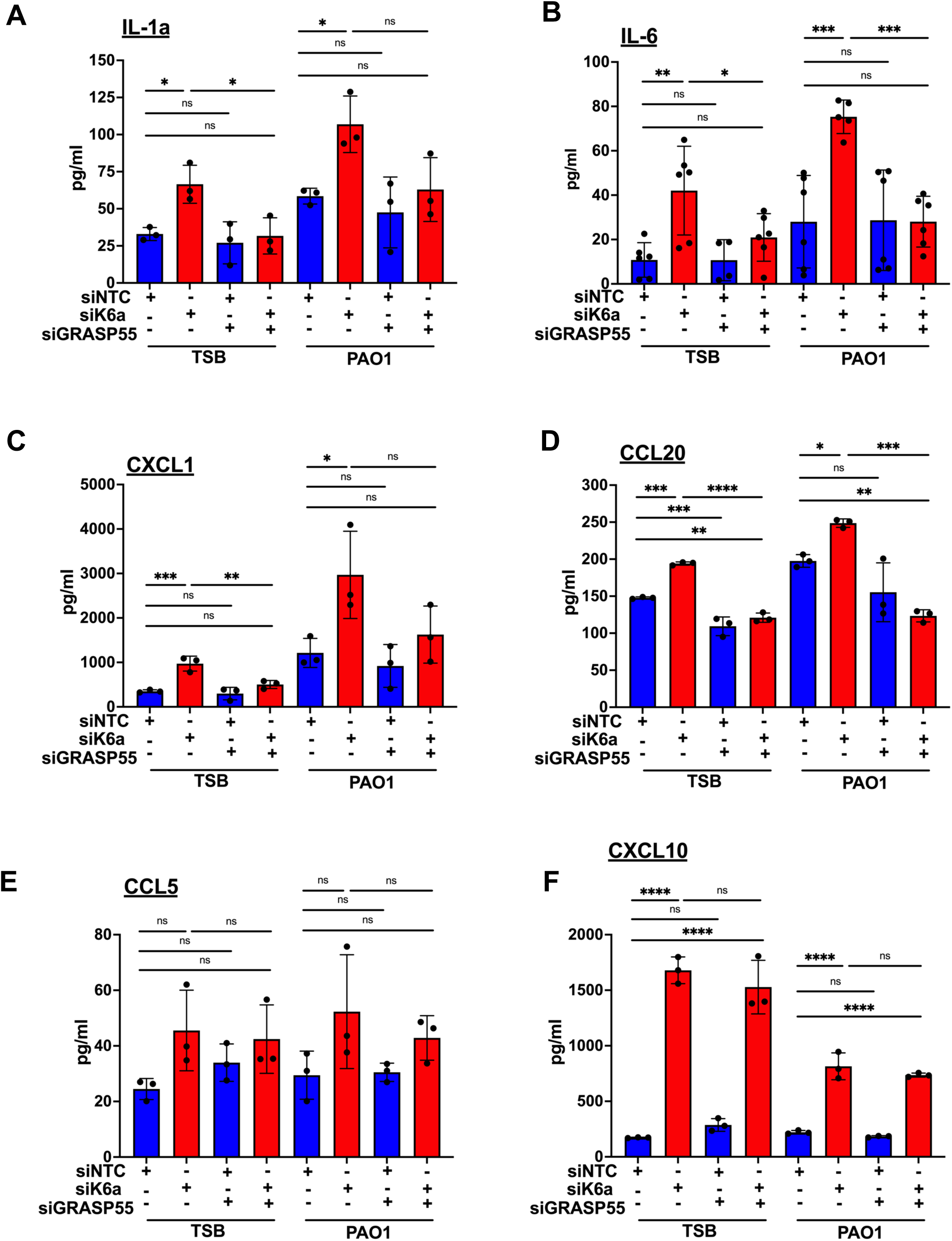
GRASP55 is required for K6a-KD-induced secretion of cytokines and chemokines. hTCEpi cells were transfected with non-targeting siRNA, or K6a– and GRASP55-specific siRNA (alone or in combination) prior to stimulation with 20% TSB or 20% sterile PAO1 culture supernatant for 18-20 h. **(A)** IL-1α, **(B)** IL-6, **(C)** CXCL1, **(D)** CCL20, **(E)** CCL5, and **(F)** CXCL10 concentrations were determined by ELISA. Data is shown as mean ± SD of 3-6 independent experiments. Statistical significance was assessed using one-way ANOVA with Sidak’s multiple comparisons test. *P < 0.05, **P < 0.01, ***P < 0.001, ****P < 0.0001, ns: P > 0.05.

### Increased autophagosome-mediated secretion of IL-8 from hTCEpi cells is dependent on Rab8a and Rab8b

Recent evidence suggested that the Rab family of small GTPases, traditionally recognized for their involvement in membrane trafficking and fusion events ^74, 75^, exerts significant influence over the autophagic process ^76^. Notably, Rab8a, a key regulator of polarized sorting towards the plasma membrane, has been reported to regulate autophagy-dependent secretion of IL-1β and HMGB1 ^34^. To examine whether Rab8a influences autophagosome-mediated IL-8 secretion from hTCEpi cells, we conducted knockdown experiments targeting Rab8a and K6a. In line with IL-1β secretion ^34^, our findings demonstrated that simultaneous depletion of Rab8a and K6a effectively nullified the augmentative effect of K6a knockdown on IL-8 secretion (Figure 7A), mirroring the observations about GRASP55. Conversely, no significant difference in IL-8 secretion was observed between WT cells and Rab8a-KD hTCEpi cells (Figure 7A). Unlike Rab8a, Rab8b is known to regulate the maturation phase of degradative autophagy by interacting with TBK-1. This interaction leads to the phosphorylation of autophagy adaptor proteins and is regarded as a potential marker for the degradative autophagy ^37, 77^. To test whether Rab8b also plays a role in excess IL-8 secretion from hTCEpi cells, we performed knockdown experiments targeting Rab8b alongside K6a. Surprisingly, akin to Rab8a depletion in K6a-KD cells, simultaneous depletion of Rab8b and K6a also abolished the augmentative effect of K6a knockdown on IL-8 secretion (Figure 7B). In conclusion, these findings highlighted the involvement of systems for polarized vesicular transport to the plasma membrane in autophagy-dependent unconventional secretion, and underscored the essential roles played by Rab8a and Rab8b in facilitating effective autophagosome-mediated IL-8 secretion from K6a-KD hTCEpi cells.

**Figure 7.**
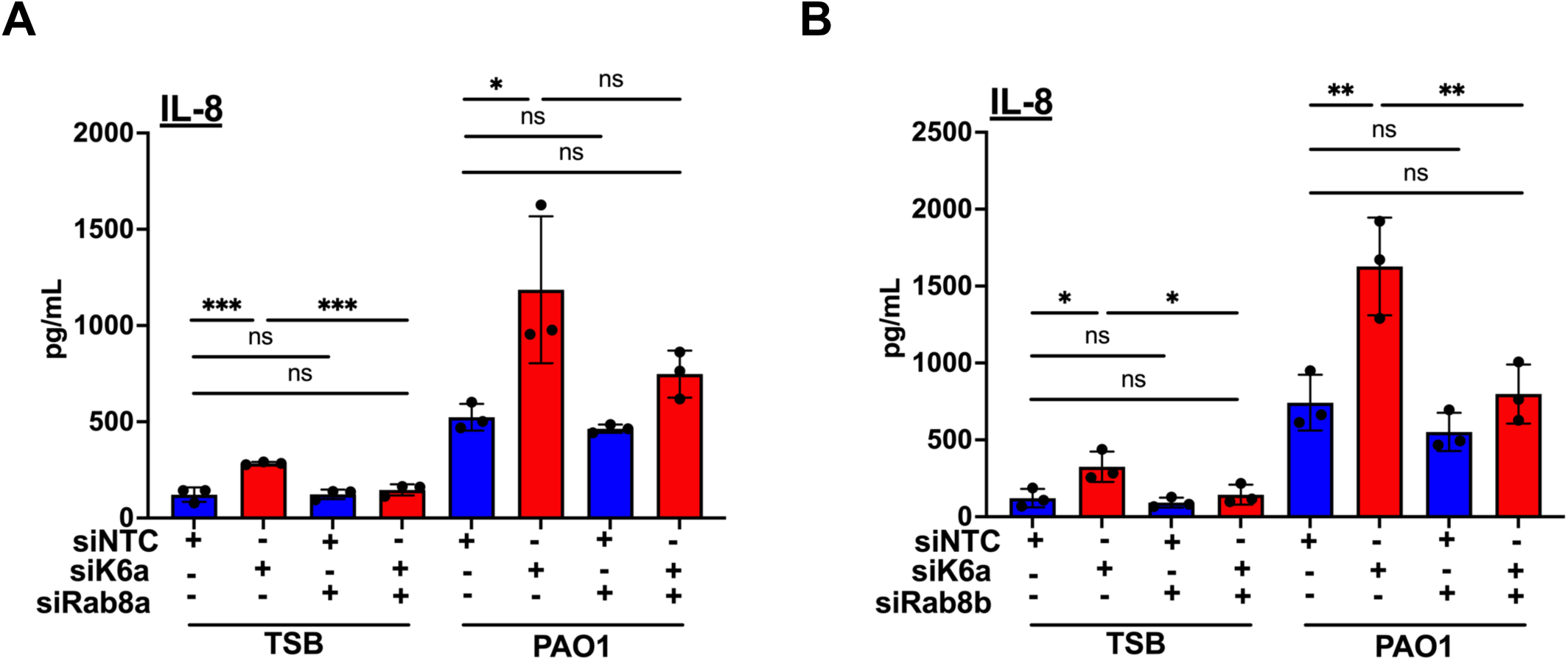
Rab8a and Rab8b are involved in K6a-KD-induced IL-8 secretion. hTCEpi cells were transfected with non-targeting siRNA, or K6a-specific siRNA alone or in combination with **(A)** Rab8a, or **(B)** Rab8b, prior to stimulation with 20% TSB or 20% sterile PAO1 culture supernatant for 18-20 h. Concentration of secreted IL-8 in culture supernatants was assessed using ELISA. Data was pooled from three independent experiments and shown as mean ± SD. Statistical significance was assessed using one-way ANOVA with Sidak’s multiple comparisons test. *P < 0.05, **P < 0.01, ***P < 0.001, ns: P > 0.05.

## DISCUSSION

In this study, we have demonstrated that knockdown of K6a in corneal epithelial cells (hTCEpi) leads to increased secretion of proinflammatory cytokines compared to WT cells. This occurs under basal condition and in response to a mixture of inflammatory ligands in bacterial culture supernatants, and our data indicated that an unconventional secretory pathway known as secretory autophagy is responsible. While degradative autophagy is known to play a crucial role in innate immunity by preventing an excessive inflammation ^78, 79^, this study highlights the contribution of secretory autophagy to exacerbated corneal inflammation, and the importance of K6a in restricting Sec16A-mediated GRASP55-dependent biogenesis of secretory autophagosomes that release proinflammatory cytokines and chemokines.

Our investigation started with multiplex assays and conventional ELISA measuring cytokine secretion from hTCEpi cells untreated or treated with cell-free bacterial culture supernatants containing a mixture of inflammatory Toll-like receptor ligands. As multiple cytokines released from K6a-KD cells are upregulated as compared to WT cells, these findings are consistent with our recent report, which has shown that K6a knockdown causes increased NF-κB activation and cytokine expression in hTCEpi cells and mouse corneal epithelium challenged with purified LPS and LTA, and that mechanistically, ELKS-interacting K6a is a newfound negative regulator of non-canonical NF-κB signaling for preventing excessive expression of proinflammatory cytokines and chemokines at the transcriptional level ^27^. In this study, K6a interactome analysis has identified Sec16A and showed various autophagy and secretory pathways enriched in hTCEpi cells; so silencing K6a has led to elevated LC3-II level (lysate protein and cytoplasmic puncta), whether or not the cells are stimulated with inflammatory ligands, suggesting a possible relationship between K6a, autophagy and cytokine secretion. A recent report has shown that keratin 8 (K8) promotes autophagosome-lysosome fusion in retinal pigment epithelial cells when subjected to oxidative stress ^23^. Additionally, proteomic data has established a connection between secretory pathways and the machinery of autophagy, revealing that Endoplasmic Reticulum Exit Sites (ERES) are fundamental components in the hierarchical assembly of the autophagy machinery ^80^. These initial findings have led us to hypothesize that K6a is involved in the regulation of corneal inflammation through its influence on autophagy and secretory pathways.

Sec16A, a conserved peripheral membrane protein localized at the ERES, has the capacity to bind nearly all COPII subunits and governs the COPII-coated vesicle dynamics ^50^. In endoplasmic reticulum (ER)-to-Golgi transport, particularly in Drosophila, Sec16A plays an indispensable role as its absence leads to a severe inhibition in protein exit from the ER ^81^. In this study, we discovered a negative regulatory mechanism of K6a; specifically, the interaction between K6a and Sec16A effectively suppresses Sec16A-dependent IL-8 release from hTCEpi cells. It is noteworthy that chronic increase in cargo load for ER export can induce an adaptive response to support the increased capacity of secretory flux. This adaptive response involves Sec16A recruitment from the cytosol to endoplasmic reticulum exit sites (ERES) to increase the formation of new ERES and COPII vesicles ^82^. In the case of K6a-KD hTCEpi cells when hypersecretion of cytokines may resemble chronic overload of cargo, we have observed a substantial increase in Sec16A immunofluorescence signal appearing as localized puncta within the cells, presumably at ERES. Corroborating with emerging evidence demonstrating the role of Sec16A in secretory autophagy in mammals and yeasts ^41, 83^, we have further shown that LC3-II^+^ autophagosome formation enhanced by K6a knockdown is abrogated by simultaneous knockdown of Sec16A, suggesting that Sec16A being free from restraint in K6a-KD cells is critical to the process of secretory autophagosome formation originating at the ERES ^69, 70^. Specifically, our findings highlight the importance of K6a in regulating recruitment of Sec16A to ERES, where vesicle formation for ER cargo exit appears to be crucial for subsequent autophagosome formation involved in unconventional secretion of excess IL-8 from hTCEpi cells under basal and inflammatory conditions.

Although GRASP55 was initially characterized as Golgi reassembly and stacking protein, later studies indicated that it is also involved in unconventional trafficking of cytosolic and transmembrane proteins ^32, 71, 84–87^. Notably, evidence supporting GRASP55’s involvement in unconventional secretion of inflammatory mediators is mounting ^34, 40, 42, 88^ while the mechanisms remain to be elucidated. In K6a-KD human corneal epithelial cells, we demonstrated that GRASP55 is associated with non-degradative autophagosomes and required for their formation, and that knockdown of GRASP55 reverses the augmentative effect of K6a-KD on secretion of cytokines and chemokines, specifically IL-1α, IL-6, IL-8, CXCL1 and CCL20. These findings highlight the functional role of GRASP55 in unconventional protein secretion as the positive regulator of secretory autophagosome formation. This is different from a previous report in which GRASP55 was found to mediate autophagy-dependent secretion of mutant huntingtin by promoting autophagosome-lysosome fusion (autophagic flux) ^71^.

Several studies have provided evidence on Rab8’s direct participation in unconventional protein secretion. For instance, Rab8a’s role has been demonstrated in unconventional secretion of ANXA2, insulin-degrading enzyme (IDE), and lysozyme ^89^. In addition, Rab8a is required for autophagy-dependent secretion of leaderless cytokine interleukin-1β (IL-1β) from murine bone marrow-derived macrophages ^34^. In our study, we tested two different Rab8 isoforms Rab8a and Rab8b, known for their role in autophagosome maturation ^37, 77^. Our findings demonstrated that both Rab8a and Rab8b proteins played a similar role in IL-8 release from K6a-KD hTCEpi cells, potentially indicating a context and tissue-specific function for these distinct Rab8a and Rab8b isoforms.

In summary, this investigation has, for the first time, demonstrated the involvement of cytokeratin K6a in autophagy-dependent inhibition of inflammation at the ocular surface. We believe that K6a impedes the function of a crucial secretory protein, Sec16A, leading to the suppression of secretory autophagosome formation and the subsequent GRASP55-dependent release of proinflammatory mediators. This study provides a novel perspective on the intrinsic cellular mechanisms that regulate inflammation at epithelial barriers both in basal as well as in response to inflammation induction.

## ACKNOWLEDGEMENT

We thank the LRI Electron Microscopy Core for EM support and services. This work was supported by NIH/NEI grants R01EY023000 and R01EY030577 (C.T.), NIH/NEI Core Grant P30EY025585 (Cole Eye Institute), and research grants from Research to Prevent Blindness and Cleveland Eye Bank (Cole Eye Institute). The Fusion Lumos instrument was purchased via an NIH shared instrument grant, 1S10OD023436-01.

## FIGURE LEGENDS

**Figure S1**. **Transmission electron microscopy revealed a higher number of autophagic vesicles in K6a-KD cells.** hTCEpi cells were seeded in a 4-well chamber slide and K6a was knocked down by siRNA. Non-targeting siRNA (siNTC) served as control. Three days post-knockdown, cells were treated with 20% TSB or 20% PAO1 culture supernatant for 24 h. **(A, C)** Cells were fixed and imaged by TEM. N, nucleus; Ap, autophagic vesicle. **(B, D)** Autophagic vesicles per cell were quantified for 3-5 cells. t-test was used to determine significance. *P < 0.05, **P < 0.01, ***P < 0.001, ****P < 0.0001, “ns” denotes non-significant results (P > 0.05).

**Figure S2**. **Immuno-transmission electron microscopy demonstrated the co-occurrence of LC3-II and GRASP55 within autophagosomes.** hTCEpi cells were subjected to siRNA knockdown of K6a with non-targeting siNTC as control. After 3 days, cells were exposed to either 20% TSB or 20% PAO1 culture supernatant for 18 h. Cells were fixed and incubated with anti-LC3 and anti-GRASP55 antibodies, followed by 5 nm (black filled arrow) and 15 nm (hollow arrow) gold-labeled secondary antibodies, respectively. “AP” denotes autophagosome. Scale bar = 200nm.

## Notes

### Competing Interest Statement

The authors have declared no competing interest.

## REFERENCES

1. Bourcier T, Thomas F, Borderie V, Chaumeil C, Laroche L. Bacterial keratitis: predisposing factors, clinical and microbiological review of 300 cases. British Journal of Ophthalmology. 2003;87(7):834–8.

2. Gayton JL. Etiology, prevalence, and treatment of dry eye disease. Clinical ophthalmology. 2009:405–12.

3. Wang B, Yang S, Zhai H-L, Zhang Y-Y, Cui C-X, Wang J-Y, Xie L-X. A comparative study of risk factors for corneal infection in diabetic and non-diabetic patients. International journal of ophthalmology. 2018;11(1):43.

4. Cabrera-Aguas M, Khoo P, Watson SL. Infectious keratitis: A review. Clinical & Experimental Ophthalmology. 2022;50(5):543–62.

5. Ray KJ, Srinivasan M, Mascarenhas J, Rajaraman R, Ravindran M, Glidden DV, Oldenburg CE, Sun CQ, Zegans ME, McLeod SD. Early addition of topical corticosteroids in the treatment of bacterial keratitis. JAMA ophthalmology. 2014;132(6):737–41.

6. Hazlef LD, Hendricks RL. Reviews for immune privilege in the year 2010: immune privilege and infection. Ocular immunology and inflammation. 2010;18(4):237–43.

7. Ueta M, Kinoshita S. Ocular surface inflammation mediated by innate immunity. Eye & contact lens. 2010;36(5):269–81.

8. Liu J, Li Z. Resident innate immune cells in the cornea. Frontiers in immunology. 2021;12:620284.

9. Nestle FO, Di Meglio P, Qin J-Z, Nickoloff BJ. Skin immune sentinels in health and disease. Nature Reviews Immunology. 2009;9(10):679–91.

10. Hewif RJ, Lloyd CM. Regulation of immune responses by the airway epithelial cell landscape. Nat Rev Immunol. 2021;21(6):347–62. Epub 20210113. doi: 10.1038/s41577-020-00477-9. PubMed PMID: 33442032; PMCID: PMC7804588.

11. Hooper LV. Epithelial cell contributions to intestinal immunity. Adv Immunol. 2015;126:129–72. Epub 20150207. doi: 10.1016/bs.ai.2014.11.003. PubMed PMID: 25727289.

12. Suzuki K, Saito J, Yanai R, Yamada N, Chikama T-i, Seki K, Nishida T. Cell–matrix and cell– cell interactions during corneal epithelial wound healing. Progress in retinal and eye research. 2003;22(2):113–33.

13. Sugrue SP, Zieske JD. ZO1 in corneal epithelium: association to the zonula occludens and adherens junctions. Experimental eye research. 1997;64(1):11–20.

14. Govindarajan B, Gipson IK. Membrane-tethered mucins have multiple functions on the ocular surface. Experimental eye research. 2010;90(6):655–63.

15. Pearlman E, Sun Y, Roy S, Karmakar M, Hise AG, Szczotka-Flynn L, Ghannoum M, Chinnery HR, McMenamin PG, Rietsch A. Host defense at the ocular surface. International reviews of immunology. 2013;32(1):4–18.

16. Tam C, Mun JJ, Evans DJ, Fleiszig SM. Cytokeratins mediate epithelial innate defense through their antimicrobial properties. The Journal of clinical investigation. 2012;122(10):3665–77.

17. Takahashi K, Paladini RD, Coulombe PA. Cloning and characterization of multiple human genes and cDNAs encoding highly related type II keratin 6 isoforms. Journal of Biological Chemistry. 1995;270(31):18581–92.

18. Dyrlund TF, Poulsen ET, Scavenius C, Nikolajsen CL, Thøgersen IB, Vorum H, Enghild JJ. Human cornea proteome: identification and quantitation of the proteins of the three main layers including epithelium, stroma, and endothelium. Journal of proteome research. 2012;11(8):4231–9.

19. Smith FJ, Liao H, Cassidy AJ, Stewart A, Hamill KJ, Wood P, Joval I, Van Steensel MA, Björck E, Callif-Daley F, editors. The genetic basis of pachyonychia congenita. Journal of Investigative Dermatology Symposium Proceedings; 2005: Elsevier.

20. Lehmann S, Leube R, Schwarz N. Keratin 6a mutations lead to impaired mitochondrial quality control. British Journal of Dermatology. 2020;182(3):636–47.

21. Dong X-M, Liu E-D, Meng Y-X, Liu C, Bi Y-L, Wu H-W, Jin Y-C, Yao J-H, Tang L-J, Wang J. Keratin 8 limits TLR-triggered inflammatory responses through inhibiting TRAF6 polyubiquitination. Scientific reports. 2016;6(1):32710.

22. Roth W, Kumar V, Beer H-D, Richter M, Wohlenberg C, Reuter U, Thiering S, Staratschek-Jox A, Hofmann A, Kreusch F. Keratin 1 maintains skin integrity and participates in an inflammatory network in skin through interleukin-18. Journal of cell science. 2012;125(22):5269–79.

23. Baek A, Yoon S, Kim J, Baek YM, Park H, Lim D, Chung H, Kim DE. Autophagy and KRT8/keratin 8 protect degeneration of retinal pigment epithelium under oxidative stress. Autophagy. 2017;13(2):248–63. Epub 20170103. doi: 10.1080/15548627.2016.1256932. PubMed PMID: 28045574; PMCID: PMC5324842.

24. Chan JK, Yuen D, Too PH-M, Sun Y, Willard B, Man D, Tam C. Keratin 6a reorganization for ubiquitin–proteasomal processing is a direct antimicrobial response. Journal of Cell Biology. 2018;217(2):731–44.

25. Lee JT, Wang G, Tam YT, Tam C. Membrane-Active Epithelial Keratin 6A Fragments (KAMPs) Are Unique Human Antimicrobial Peptides with a Non-alphabeta Structure. Front Microbiol. 2016;7:1799. Epub 20161111. doi: 10.3389/fmicb.2016.01799. PubMed PMID: 27891122; PMCID: PMC5105358.

26. Sun Y, Chan J, Bose K, Tam C. Simultaneous control of infection and inflammation with keratin-derived antibacterial peptides targeting TLRs and co-receptors. Science Translational Medicine. 2023;15(686):eade2909.

27. Chan JK, Sun Y, Bhushan A, Willard B, Tam C. Keratin 6a afenuates Toll-like receptor-triggered proinflammatory response in corneal epithelial cells by suppressing ELKS/IKKɛ-dependent activation of NF-κB. bioRxiv. 2023:2023.10. 24.563888.

28. Miller EA, Beilharz TH, Malkus PN, Lee MC, Hamamoto S, Orci L, Schekman R. Multiple cargo binding sites on the COPII subunit Sec24p ensure capture of diverse membrane proteins into transport vesicles. Cell. 2003;114(4):497–509.

29. Ofe S, Barlowe C. Sorting signals can direct receptor-mediated export of soluble proteins into COPII vesicles. Nature Cell Biology. 2004;6(12):1189–94.

30. Nickel W. The mystery of nonclassical protein secretion: a current view on cargo proteins and potential export routes. European Journal of Biochemistry. 2003;270(10):2109–19.

31. Giuliani F, Grieve A, Rabouille C. Unconventional secretion: a stress on GRASP. Current opinion in cell biology. 2011;23(4):498–504.

32. Nickel W, Rabouille C. Mechanisms of regulated unconventional protein secretion. Nature reviews Molecular cell biology. 2009;10(2):148–55.

33. Bruns C, McCaffery JM, Curwin AJ, Duran JM, Malhotra V. Biogenesis of a novel compartment for autophagosome-mediated unconventional protein secretion. Journal of Cell Biology. 2011;195(6):979–92.

34. Dupont N, Jiang S, Pilli M, Ornatowski W, Bhafacharya D, Deretic V. Autophagy-based unconventional secretory pathway for extracellular delivery of IL-1β. The EMBO journal. 2011;30(23):4701–11.

35. Duran JM, Kinseth M, Bossard C, Rose DW, Polishchuk R, Wu CC, Yates J, Zimmerman T, Malhotra V. The role of GRASP55 in Golgi fragmentation and entry of cells into mitosis. Molecular biology of the cell. 2008;19(6):2579–87.

36. Manjithaya R, Anjard C, Loomis WF, Subramani S. Unconventional secretion of Pichia pastoris Acb1 is dependent on GRASP protein, peroxisomal functions, and autophagosome formation. Journal of Cell Biology. 2010;188(4):537–46.

37. Jiang S, Dupont N, Castillo EF, Deretic V. Secretory versus degradative autophagy: unconventional secretion of inflammatory mediators. Journal of innate immunity. 2013;5(5):471–9.

38. Murai H, Okazaki S, Hayashi H, Kawakita A, Hosoki K, Yasutomi M, Sur S, Ohshima Y. Alternaria extract activates autophagy that induces IL-18 release from airway epithelial cells. Biochemical and biophysical research communications. 2015;464(4):969–74.

39. Gee HY, Noh SH, Tang BL, Kim KH, Lee MG. Rescue of ΔF508-CFTR trafficking via a GRASP-dependent unconventional secretion pathway. Cell. 2011;146(5):746–60.

40. Nüchel J, Ghatak S, Zuk AV, Illerhaus A, Mörgelin M, Schönborn K, Blumbach K, Wickström SA, Krieg T, Sengle G. TGFB1 is secreted through an unconventional pathway dependent on the autophagic machinery and cytoskeletal regulators. Autophagy. 2018;14(3):465–86.

41. Piao H, Kim J, Noh SH, Kweon H-S, Kim JY, Lee MG. Sec16A is critical for both conventional and unconventional secretion of CFTR. Scientific reports. 2017;7(1):39887.

42. Zhang M, Kenny SJ, Ge L, Xu K, Schekman R. Translocation of interleukin-1β into a vesicle intermediate in autophagy-mediated secretion. elife. 2015;4:e11205.

43. Davis S, Wang J, Ferro-Novick S. Crosstalk between the secretory and autophagy pathways regulates autophagosome formation. Developmental Cell. 2017;41(1):23–32.

44. Cavalli G, Cenci S. Autophagy and protein secretion. Journal of molecular biology. 2020;432(8):2525–45.

45. Ponpuak M, Mandell MA, Kimura T, Chauhan S, Cleyrat C, Deretic V. Secretory autophagy. Current opinion in cell biology. 2015;35:106–16.

46. Keller M, Rüegg A, Werner S, Beer H-D. Active caspase-1 is a regulator of unconventional protein secretion. Cell. 2008;132(5):818–31.

47. Robertson DM, Li L, Fisher S, Pearce VP, Shay JW, Wright WE, Cavanagh HD, Jester JV. Characterization of growth and differentiation in a telomerase-immortalized human corneal epithelial cell line. Invest Ophthalmol Vis Sci. 2005;46(2):470–8. doi: 10.1167/iovs.04-0528. PubMed PMID: 15671271.

48. Fabregat A, Sidiropoulos K, Viteri G, Forner O, Marin-Garcia P, Arnau V, D’Eustachio P, Stein L, Hermjakob H. Reactome pathway analysis: a high-performance in-memory approach. BMC Bioinformatics. 2017;18(1):142. Epub 20170302. doi: 10.1186/s12859-017-1559-2. PubMed PMID: 28249561; PMCID: PMC5333408.

49. Deretic V, Kimura T, Timmins G, Moseley P, Chauhan S, Mandell M. Immunologic manifestations of autophagy. The Journal of clinical investigation. 2015;125(1):75–84.

50. Sprangers J, Rabouille C. SEC16 in COPII coat dynamics at ER exit sites. Portland Press Ltd.; 2015.

51. Watson P, Townley AK, Koka P, Palmer KJ, Stephens DJ. Sec16 defines endoplasmic reticulum exit sites and is required for secretory cargo export in mammalian cells. Traffic. 2006;7(12):1678–87.

52. Sanchez-Wandelmer J, KBstakis NT, Reggiori F. ERES: sites for autophagosome biogenesis and maturation? Journal of cell science. 2015;128(2):185–92.

53. Bao J, Huang M, Petranovic D, Nielsen J. Moderate expression of SEC16 increases protein secretion by Saccharomyces cerevisiae. Applied and environmental microbiology. 2017;83(14):e03400–16.

54. Noh SH, Gee HY, Kim Y, Piao H, Kim J, Kang CM, Lee G, Mook-Jung I, Lee Y, Cho JW. Specific autophagy and ESCRT components participate in the unconventional secretion of CFTR. Autophagy. 2018;14(10):1761–78.

55. Wang AL, Boulton ME, Dunn J, William A, Rao HV, Cai J, Lukas TJ, Neufeld AH. Using LC3 to monitor autophagy flux in the retinal pigment epithelium. Autophagy. 2009;5(8):1190–3.

56. Mizushima N, Yoshimori T, Levine B. Methods in mammalian autophagy research. Cell. 2010;140(3):313–26.

57. Klinonsky D. Guidelines for the use and interpretation of assays for monitoring autophagy. Autophagy. 2016.

58. Jung M, Choi H, Mun JY. The autophagy research in electron microscopy. Appl Microsc. 2019;49(1):11. Epub 20191106. doi: 10.1186/s42649-019-0012-6. PubMed PMID: 33580401; PMCID: PMC7809580.

59. Mizushima N, Yoshimori T. How to interpret LC3 immunobloyng. Autophagy. 2007;3(6):542–5.

60. Giménez-Xavier P, Francisco R, Platini F, Pérez R, Ambrosio S. LC3-I conversion to LC3-II does not necessarily result in complete autophagy. International journal of molecular medicine. 2008;22(6):781–5.

61. Mauvezin C, Nagy P, Juhasz G, Neufeld TP. Autophagosome-lysosome fusion is independent of V-ATPase-mediated acidification. Nat Commun. 2015;6:7007. Epub 20150511. doi: 10.1038/ncomms8007. PubMed PMID: 25959678; PMCID: PMC4428688.

62. Bjørkøy G, Lamark T, Brech A, Outzen H, Perander M, Øvervatn A, Stenmark H, Johansen T. p62/SQSTM1 forms protein aggregates degraded by autophagy and has a protective effect on huntingtin-induced cell death. The Journal of cell biology. 2005;171(4):603–14.

63. Bjørkøy G, Lamark T, Pankiv S, Øvervatn A, Brech A, Johansen T. Monitoring autophagic degradation of p62/SQSTM1. Methods in enzymology. 2009;452:181–97.

64. Kimura S, Noda T, Yoshimori T. Dissection of the autophagosome maturation process by a novel reporter protein, tandem fluorescent-tagged LC3. Autophagy. 2007;3(5):452–60.

65. Narita M, Young AR, Arakawa S, Samarajiwa SA, Nakashima T, Yoshida S, Hong S, Berry LS, Reichelt S, Ferreira M. Spatial coupling of mTOR and autophagy augments secretory phenotypes. Science. 2011;332(6032):966–70.

66. Kraya AA, Piao S, Xu X, Zhang G, Herlyn M, Gimofy P, Levine B, Amaravadi RK, Speicher DW. Identification of secreted proteins that reflect autophagy dynamics within tumor cells. Autophagy. 2015;11(1):60–74.

67. Codogno P, Meijer AJ. Atg5: more than an autophagy factor. Nature cell biology. 2006;8(10):1045–7.

68. Thorburn J, Horita H, Redzic J, Hansen K, Frankel AE, Thorburn A. Autophagy regulates selective HMGB1 release in tumor cells that are destined to die. Cell Death & Differentiation. 2009;16(1):175–83.

69. Ge L, Melville D, Zhang M, Schekman R. The ER–Golgi intermediate compartment is a key membrane source for the LC3 lipidation step of autophagosome biogenesis. elife. 2013;2:e00947.

70. Curwin AJ, Brouwers N, Alonso Y, Adell M, Teis D, Turacchio G, Parashuraman S, Ronchi P, Malhotra V. ESCRT-III drives the final stages of CUPS maturation for unconventional protein secretion. Elife. 2016;5:e16299.

71. Ahat E, Bui S, Zhang J, da Veiga Leprevost F, Sharkey L, Reid W, Nesvizhskii AI, Paulson HL, Wang Y. GRASP55 regulates the unconventional secretion and aggregation of mutant huntingtin. Journal of Biological Chemistry. 2022;298(8).

72. Kim J, Noh SH, Piao H, Kim DH, Kim K, Cha JS, Chung WY, Cho HS, Kim JY, Lee MG. Monomerization and ER relocalization of GRASP is a requisite for unconventional secretion of CFTR. Traffic. 2016;17(7):733–53.

73. Nüchel J, Tauber M, Nolte JL, Mörgelin M, Türk C, Eckes B, Demetriades C, Plomann M. An mTORC1-GRASP55 signaling axis controls unconventional secretion to reshape the extracellular proteome upon stress. Molecular Cell. 2021;81(16):3275–93. e12.

74. Homma Y, Hiragi S, Fukuda M. Rab family of small GTPases: an updated view on their regulation and functions. The FEBS journal. 2021;288(1):36–55.

75. Wandinger-Ness A, Zerial M. Rab proteins and the compartmentalization of the endosomal system. Cold Spring Harbor perspectives in biology. 2014;6(11):a022616.

76. Bento CF, Puri C, Moreau K, Rubinsztein DC. The role of membrane-trafficking small GTPases in the regulation of autophagy. Journal of cell science. 2013;126(5):1059–69.

77. Pilli M, Arko-Mensah J, Ponpuak M, Roberts E, Master S, Mandell MA, Dupont N, Ornatowski W, Jiang S, Bradfute SB. TBK-1 promotes autophagy-mediated antimicrobial defense by controlling autophagosome maturation. Immunity. 2012;37(2):223–34.

78. Nakahira K, Haspel JA, Rathinam VA, Lee S-J, Dolinay T, Lam HC, Englert JA, Rabinovitch M, Cernadas M, Kim HP. Autophagy proteins regulate innate immune responses by inhibiting the release of mitochondrial DNA mediated by the NALP3 inflammasome. Nature immunology. 2011;12(3):222–30.

79. Castillo EF, Dekonenko A, Arko-Mensah J, Mandell MA, Dupont N, Jiang S, Delgado-Vargas M, Timmins GS, Bhafacharya D, Yang H. Autophagy protects against active tuberculosis by suppressing bacterial burden and inflammation. Proceedings of the National Academy of Sciences. 2012;109(46):E3168–E76.

80. Graef M, Friedman JR, Graham C, Babu M, Nunnari J. ER exit sites are physical and functional core autophagosome biogenesis components. Molecular biology of the cell. 2013;24(18):2918–31.

81. Aguilera-Gomez A, Zacharogianni M, van Oorschot MM, Genau H, Grond R, Veenendaal T, Sinsimer KS, Gavis ER, Behrends C, Rabouille C. Phospho-Rasputin stabilization by Sec16 is required for stress granule formation upon amino acid starvation. Cell reports. 2017;20(4):935–48.

82. Farhan H, Weiss M, Tani K, Kaufman RJ, Hauri HP. Adaptation of endoplasmic reticulum exit sites to acute and chronic increases in cargo load. The EMBO journal. 2008;27(15):2043–54.

83. Ishihara N, Hamasaki M, Yokota S, Suzuki K, Kamada Y, Kihara A, Yoshimori T, Noda T, Ohsumi Y. Autophagosome requires specific early Sec proteins for its formation and NSF/SNARE for vacuolar fusion. Molecular biology of the cell. 2001;12(11):3690–702.

84. Malhotra V. Unconventional protein secretion: an evolving mechanism. The EMBO journal. 2013;32(12):1660–4.

85. Rabouille C. Pathways of unconventional protein secretion. Trends in cell biology. 2017;27(3):230–40.

86. Gee HY, Kim J, Lee MG, editors. Unconventional secretion of transmembrane proteins. Seminars in cell & developmental biology; 2018: Elsevier.

87. Zhang X, Wang Y. Nonredundant Roles of GRASP55 and GRASP65 in the Golgi Apparatus and Beyond. Trends in biochemical sciences. 2020;45(12):1065–79.

88. Kim YH, Kwak MS, Lee B, Shin JM, Aum S, Park IH, Lee MG, Shin J-S. Secretory autophagy machinery and vesicular trafficking are involved in HMGB1 secretion. Autophagy. 2021;17(9):2345–62.

89. Claude-Taupin A, Jia J, Mudd M, Deretic V. Autophagy’s secret life: secretion instead of degradation. Essays in Biochemistry. 2017;61(6):637–47.

