## Supplementary figures and images for "Epithelial cytokeratin 6a restricts secretory autophagy of proinflammatory cytokines by interacting with Sec16A"

### Supplemental Figure 1 and 2

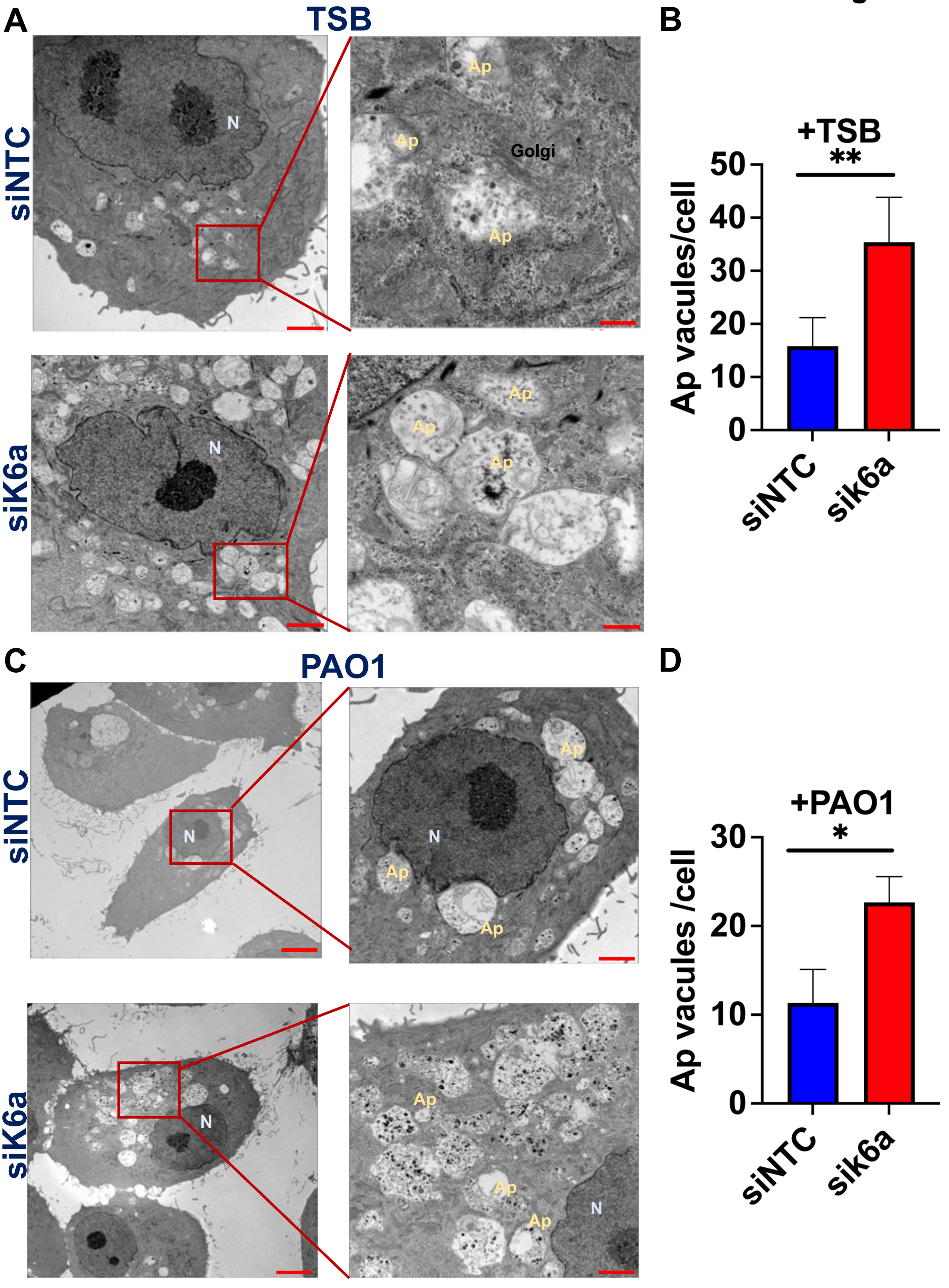

**Figure S2**

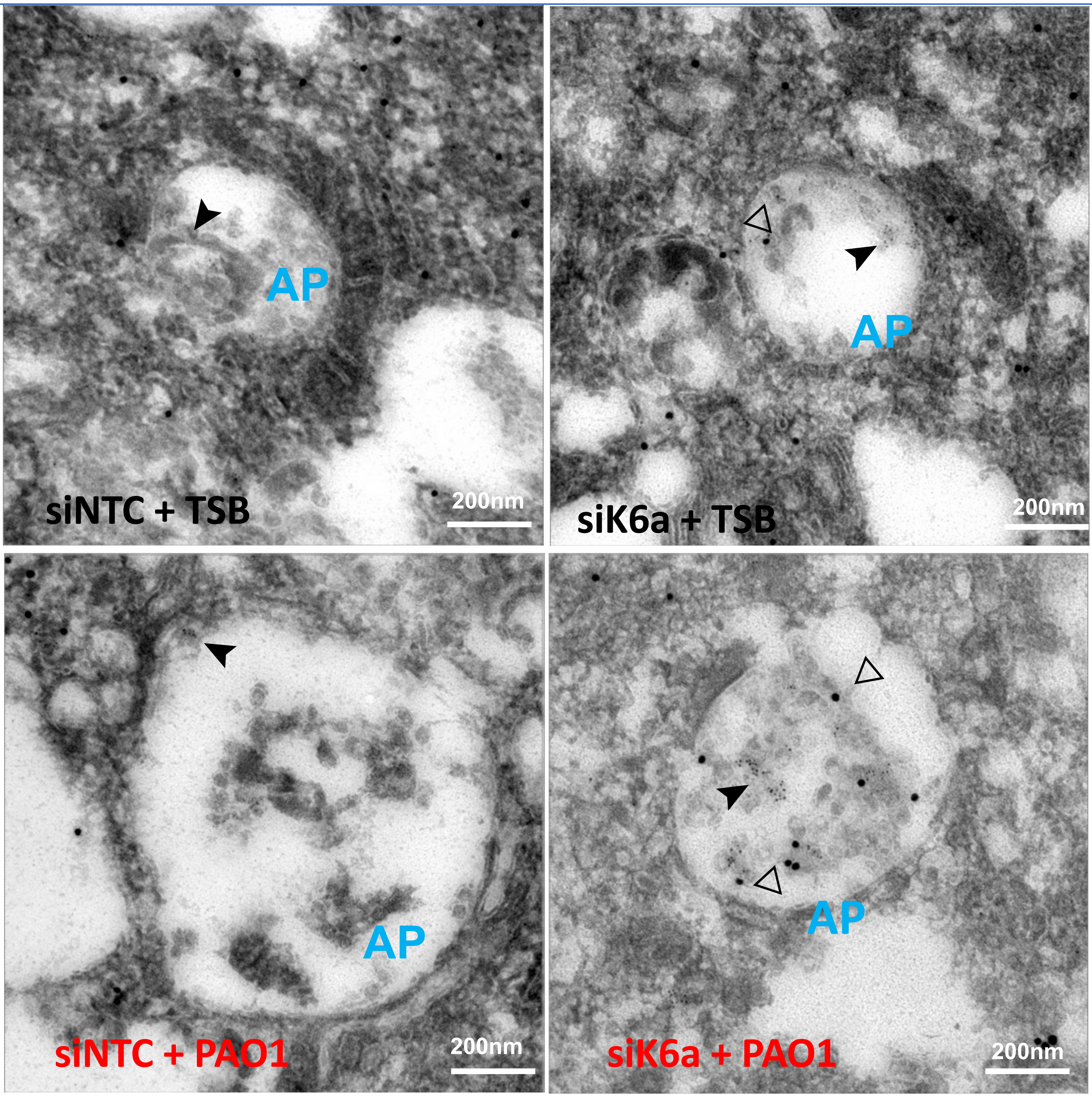


**LC3-II (5 nm, ➤); GRASP55 (15 nm, ▷)**

**AP: Autophagosome**
